# Serological proteomic screening and evaluation of a recombinant egg antigen for the diagnosis of low-intensity infections in endemic area in Brazil Antigen target in schistosomiasis diagnosis

**DOI:** 10.1101/463059

**Authors:** Vanessa Silva-Moraes, Lisa M Shollenberger, William Castro-Borges, Ana Lucia Teles Rabello, Donald A Harn, Lia Carolina Soares Medeiros, Wander de Jesus Jeremias, Liliane Maria Vidal Siqueira, Caroline Stephane Salviano Pereira, Maria Luysa Camargos Pedrosa, Nathalie Bonatti Franco Almeida, Aureo Almeida, Jose Roberto Lambertucci, Nídia Francisca de Figueiredo Carneiro, Paulo Marcos Zech Coelho, Rafaella Fortini Grenfell e Queiroz

## Abstract

**Background:** Despite decades of employment of control programs, schistosomiasis remains a global public health problem. To further reduce prevalence and intensity of infection, or to achieve the goal of elimination in low-endemic areas, there need to be better diagnostic tools to detect low intensity infections in low-endemic areas as Brazil. The rationale for development of new diagnostic tools is because in low-endemic settings, the standard Kato-Katz diagnostic method loses its sensitivity and misses low intensity infections. To develop new diagnostic tools, we employed a proteomics approach search for biomarkers associated with schistosome-specific immune responses to develop sensitive and specific new methods for immunodiagnosis.

**Methods and findings:** Immunoproteomic analysis was performed on an egg extract of *Schistosoma mansoni* using pooled sera from infected or non-infected individuals from a low-endemic area of Brazil. Cross reactivity with other helminth parasites was determined using pooled sera from individuals infected with different parasitic helminths. Using this approach we identified 23 spots recognized by schistosome acute and chronic sera samples. To identify immunoreactive spots that are likely glycan epitopes, we compared immunoreactivity of spots treated by sodium metaperiodate oxidation of egg extract. This treatment yielded 12/23 spots maintaining immunoreactivity, suggesting they are protein epitopes. From these 12 spots, 11 spots cross-reacted with sera from infection with other helminths and 10 spots cross-reacted with the negative control group. Spot number 5 was exclusively immunoreactive with sera from schistosome-infected groups in native and deglycosylated conditions and corresponds to a Major Egg Antigen. We expressed the Major egg antigen as a recombinant protein and showed by western blot a similar recognition pattern of that of the native protein. IgG-ELISA gave a sensitivity of 87.10% and specificity of 89.09% represented by area under ROC curve of 0.95. IgG-ELISA performed better than the conventional K-K (2 slides) identifying 56/64 cases harboring 1-10 egg per gram of feces that were undiagnosed by K-K parasitological technique.

**Conclusions:** The serological proteome approach was able to identify a new diagnostic candidate. The recombinant egg antigen provided good performance in IgG-ELISA to detect individuals with extreme low-intensity infections (1 egg per gram of feces). Therefore, the IgG-ELISA using this newly identified recombinant major egg antigen can be a useful tool to be combined with other techniques in low-endemic areas to determine the true prevalence of schistosome infection that is underestimated by Kato-Katz method. Further, to overcome the complexity of ELISA in the field, a second-generation of antibody-based rapid diagnostic tests (RDT) can be developed.

**Author Summary:** Schistosomiasis remains a serious global public health problem. Detecting parasite eggs in patient stool samples using the Kato-Katz (KK) method is the standard diagnostic recommended by World Health Organization (WHO) for infection by *Schistosoma mansoni*. As a result of intensive control strategies, many previously high-endemic areas are now considered low-endemic areas and the K-K method does not function well in low-endemic areas and therefore cannot be considered the gold standard. Thus, a new emphasis on strategies to accurately diagnose low-intensity infections was outlined in plan from WHO focusing on elimination of disease as a public health problem. Successful diagnoses and treatment of the majority of infected individuals may result in elimination of a sufficient number of low-burden transmitters and consequently, in the interruption of the disease transmission. In this regard, immunological techniques have proven to be more sensitive and promising for identifying infection in low-intensity of infection positive individuals with negative K-K results. The identification of antigens is the initial step for developing new immunodiagnostic assays. In this study, we used sets of pooled human sera samples from controls to acute and chronic infections to narrow down the number of new candidate antigens via proteomic screening. Using these approaches we initially identified 12 different egg proteins in schistosome-infected individuals (acute and chronic phase). A single antigen identified as Major Egg Antigen was shown to be highly specific as this antigen was not recognized by sera from negative or patients infected with other helminths. The recombinant Major Egg Antigen protein functioned in ELISA as a highly sensitive and specific antigen to detect patient IgG-antibodies. Recombinant Major Egg Antigen performed significantly better to detect low-intensity infections (1 egg per gram of feces) than the K-K method using 2 slides. Therefore, using a proteomic screening approach we were able to identify a potential new candidate antigen for development of far more sensitive diagnostic assays. Further diagnostic assays employing the Major Egg Antigen could be a useful tool on its own or in combination with other methods for diagnosis of schistosome infection in populations living in extreme low-intensity endemic areas of Brazil.

## Introduction

Despite adoption of control programs, schistosomiasis is still a global public health problem. The Global Health Estimates attributed 3.51 million disability-adjusted life years (DALYs) and 10,100 disease deaths to schistosomiasis [1]. Estimates suggest that approximately 249 million people are infected with schistosomes in 78 countries around the world with more than 779 million living in endemic areas [2, 3]. The disease afflicts low-income populations in tropical and subtropical regions with varying levels of morbidity and mortality and additionally, has a significant socioeconomic impact. Brazil has the highest burden of disease in the Americas and infection is caused by *Schistosoma mansoni* [4].

Socioeconomic development, including access to basic sanitation and drug treatment with Praziquantel (PZQ), has led to a considerable reduction in the number of people infected. These same aspects have led to a reduction in morbidity, as well as to interruption of transmission in countries [5]. In Brazil, the National Schistosomiasis and Soil-transmitted Helminth Infection Survey (INPEG), conducted from 2010 to 2015, estimated that the prevalence of infection was 1.79% [6]. Despite this significant decrease after nearly 40 years of control, the disease continues to expand and acquire a new epidemiological profile, mainly in the Northeast and Southeast regions of the country [6]. Currently, Brazil has multiple endemic areas where chronically infected patients have low-intensity infections (number of eggs per gram of feces, epg, <100). In addition, Brazil has acute infection cases as a result of internal migration of infected individuals [6, 7].

PZQ mass drug administration is not conducted in Brazil. The main strategy to control and eliminate the disease is diagnosis and treatment of active cases at the primary care level [4, 8]. Currently diagnosis continues to be detection of schistosome eggs in stools by microscopic examination using the Kato-Katz technique (K-K). K-K is the World Health Organization (WHO) reference for diagnosis [9]. The K-K method is low-cost and suitable for detection of high-intensity infections. However, the K-K technique has poor sensitivity for detection of low-intensity infections that are seen in residents living in low-endemic areas (<10% prevalence, <100 epg). The low-sensitivity of the K-K method results in misdiagnosis (schistosome negative) of infected individuals, who due to lack of diagnosis, continue to contribute to disease transmission and skew actual disease prevalence. Previous studies in Brazil demonstrated that prevalence of disease has been underestimated by a factor of 2-4, due to the inability of the K-K method to detect low-intensity infections [10-14].

As a result of intensive control strategies employing praziquantel (PZQ), many previously high-endemic areas are now considered low-endemic areas where due to lack of sensitivity, K-K cannot be used as the gold standard for diagnosis [15]. Therefore, new diagnostic methods need to be developed to detect low-intensity infections. The ability to accurately diagnose low-intensity infections was outlined in plans focusing on elimination of disease as a public health problem [7, 16, 17]. Methods that can accurately diagnosis the majority of individuals will contribute to the goal of elimination of low-burden transmitters and consequently, in the interruption of the disease transmission. In this regard, molecular and immunological techniques have proven to be more sensitive and promising for identifying infection in infected individuals that are negative by K-K coproscopy results [10, 13, 14, 18-20].

Significant progress has been seen in the development of antigen-based rapid diagnostic tests (RDT), as its assembly is user-friendly in the field. The immunochromatographic point-of-care (POC) test that detects circulating cathodic antigen (CCA) in urine has been commercially available since 2008 [21, 22]. Although POC-CCA has been suggested to be a suitable substitute for K-K in *S. mansoni* prevalence mapping [22-25], its performance is still debatable in low-endemic areas [26, 27]. The majority of studies validating POC-CCA were conducted in Africa, whereas few (10) studies were conducted in Brazil, which has a significantly different prevalence and morbidity profile. In contrast to Africa, most of the low-intensity infections in Brazil are denoted as < 25 epg [11, 12, 20, 27-33].

Antibody-based methods have high sensitivity in detecting low-intensity infections and are capable of identifying loads of 1 epg [14, 19, 34-39]. Their use as screening tools combined with parasitological evaluations has decreased the false-negative cases seen when the unique analysis by 2 K-K slides is applied in endemic settings. Furthermore, another useful application of these tests is the ability to detect acute infections in individuals from non-endemic areas recently exposed to schistosomiasis-endemic settings. Since antibodies to the parasite develop during the first weeks after infection, they can be detected before eggs in the feces to yield higher sensitivities. In clinical practice, positive serology in K-K negative people from non-endemic countries is usually sufficient to prescribe treatment with PZQ [40-42].

In regards to developing an antibody-based test, the choice of antigen is the most challenging. Crude antigens, such as soluble eggs antigens (SEA) and worm antigens (SWAP), are frequently used, but they can exhibit low-sensitivity and cross-reactivity with other helminthes [43, 44]. A combination of proteomic and serological analyses have served as promising experimental approaches for screening new biomarkers in the diagnostic field [45-47]. However, there is a limited number of serological-proteomic studies involving *Schistosoma spp.* and most of them are related to searching for vaccine candidates using animal models [47-52]. Only one immunoproteomic analysis related to *S. mansoni* and human samples has been performed to date, but it also focuses on the search for vaccines candidates [52].

Since the lack of effective diagnostic assays in low-endemic areas is one of the factors that contributes to transmission and there is a need for more specific biomarkers in immunodiagnostic development, in this present work, we adopted immunoproteomic analysis to identify a new antigen candidate for schistosomiasis diagnosis. We used different sets of pooled sera from acute and chronically infected patients, as well as from helminth positive, but schistosome-negative individuals, to select highly specific schistosome antigens. Antibodies against schistosome eggs have been considered useful antigens for the diagnosis of schistosomiasis [35, 42, 53]. Therefore, we screened soluble egg extracts (SEE) by two-dimensional western blotting (2D-WB). While many studies have focused on the serologic-proteomics of adult worms, we focused on egg antigens. Antigens from eggs are highly immunogenic, but less specific, due to cross reactions provided by glycan epitopes [44, 54]. To achieve higher specificity from a recombinant antigen, we compared native SEE extracts to those oxidized by Sodium Metaperiodate (SMP), to identify those antigens whose antigens were glycan-based.

We identified 23 immunoreactive spots, which resolved to 12 differently characterized proteins. One protein was uniquely recognized by schistosome-infected patients and not recognized by sera from patients infected with other helminths or negative control sera. We produced and purified a recombinant protein for this antigen developing a conventional Enzyme-linked Immunosorbent Assay to detect antigen-specific IgG (IgG-ELISA). The recombinant egg antigen showed high sensitivity for detection of low-intensity infections that were misdiagnosed by the standard K-K (2 slides) method. The IgG-ELISA showed a useful test to diagnose the hard-to-detect patients (load of 1 epg). Furthermore, this recombinant egg antigen can be developed for use in a variety of immunodiagnostic platforms, including RDT and POC to improve schistosomiasis diagnosis in the field.

## Methods

### Ethics Statement

The present study was approved by the Ethics Committee of the Research Center Rene Rachou/Fiocruz under the following number: 893.582 11/2014 and by the National Brazilian Ethical Board under the following number: 14886. Before any research activities, the local health authorities were contacted and agreed to collaborate with the researchers from the different institutions. All enrolled participants were required to sign an informed consent form. Parents or legal guardians signed the informed consent when minors were involved. When the parasitological results were positive, the relevant individuals were informed and received free oral treatment at the local health clinic. Schistosomiasis: praziquantel (40 mg/kg for adults and 60 mg/kg for children); intestinal helminths: albendazole (400 mg); protozoan parasites: metronidazole (250 mg/2x/ 5 days).

All procedures involving animals were conducted in compliance with the Manual for the Use of Animals/FIOCRUZ and approved by the Ethics Committee on the Use of Experimental Animal (CEUA – FIOCRUZ) license number LW-31/15.

### Human sera samples

#### Chronic infection patient sera

Chronic infection sera samples were obtained from a study performed in 2009-2014 in different rural communities of Montes Claros, state of Minas Gerais, Brazil. These included Pedra Preta, Tabuas and Estreito de Miralta (491 residents, 243 women/250 men, 1-94 years old). This rural region is a schistosomiasis low-endemic area with the majority of individuals holding low-intensity infections (< 100 epg). It was selected based on environmental conditions, the presence of *Biomphalaria glabrata* snails (the intermediate host of *S. mansoni*), previously reported prevalence, and low migration rate.

The diagnosis of schistosomiasis as well as the diagnosis of other intestinal helminthes was determined via parasitological examination of 2 grams of feces. Each resident provided a stool sample that was used to make 24 K-K slides (24 × 41.7 mg = 1 gram) (Helm-Test^®^, Biomanguinhos, FIOCRUZ, Brazil) [9] and 2 analyses of Saline Gradient (SG) technique (2 × 500 mg = 1 gram) [55]. Results were reported as epg of feces for both methods. Participants infected with *S. mansoni* and/or other helminths were treated with PZQ (60 mg/kg for children and 50 mg/kg for adults) and albendazole (400 mg), respectively, in single oral dose, as recommended by the Brazilian Health Ministry. All positive patients were followed up 30, 90 and 180 days post-treatment and the same baseline procedure was performed. Individuals testing positive post-treatment were retreated as needed. Individual serum samples from all participating individuals were obtained after centrifugation of blood samples at 3000g for 5 min and were maintained at −20°C until use.

For the immunoproteomic analysis, 15 sera samples were pooled for each group. Groups were: 1) individuals positive for *S. mansoni*, but not infected with other geohelminths (INF-CR), 2) individuals positive for other geohelminths (*Ascaris lumbricoides, Trichuris trichiura and Ancylostoma*), but not *S. mansoni* (HT), and 3) individuals negative for *S. mansoni* and other geohelminths at 180 days post-treatment (PT-CR). To develop IgG-ELISA, 93 positive individuals, according to INF-CR criteria, were taken (41 women/52 men, 5-80 years old) (Table 1). The average intensity of infection was 5.4 epg, calculated by the geometric mean of the number of epg. Eighty individuals from endemic areas with negative stool examination (NEG-END) at baseline were also evaluated.

**Table 1.** Positive individuals (INF-CR) from endemic area of Minas Gerais, Brazil.

| Variables | Category | Total<br>n | % |
| --- | --- | --- | --- |
| Sex | Male | 52 | 55.9 |
|  | Female | 41 | 44.1 |
|  | Total | 93 | 100.0 |
| Age | ≤ 10 | 7 | 7.5 |
|  | 11-20 | 29 | 31.2 |
|  | 21-40 | 26 | 28.0 |
|  | 41-60 | 21 | 22.6 |
|  | >60 | 10 | 10.8 |
|  | Total | 93 | 100.0 |
| Parasite load (epg) | ≤ 10 | 64 | 68.8 |
|  | 11-25 | 12 | 12.9 |
|  | 26-50 | 3 | 3.2 |
|  | 51-99 | 8 | 8.6 |
|  | ≥ 100 | 6 | 6.5 |
|  | Total | 93 | 100.0 |

epg: eggs per gram of feces

#### Acute infection patient sera

Fifty tourists (19 women/31 men, 4-75 years old) bathed in a swimming pool supplied by a brook on a country house in the outskirts of São João del Rei, a historical city in the state of Minas Gerais, Brazil from December 2009 to March 2010 as previously described [40]. Two months later, a patient was diagnosed with schistosomal myeloradiculopathy and he reported that other tourists had symptoms consistent with acute schistosomiasis. All participants submitted to clinical, laboratory, and ultrasound examinations, and the outbreak was confirmed. Authorities investigated the species of snail surrounding the area and *Biomphalaria glabrata* was the only species found. It was determined that transmission occurred because of in-migration of infected workers who were hired to build houses in the neighborhood of a country house. This caused the non-endemic area to become a new focus of transmission.

Diagnosis of acute *S. mansoni* schistosomiasis was based on epidemiologic data (recent contact with stream water in an endemic area), clinical data (i.e. cercarial dermatitis, acute enterocolitis, fever, cough, malaise, paraplegia, pulmonary involvement, hepatomegaly and/or splenomegaly), laboratory assays (i.e. eosinophilia, IgG antibodies against soluble worm antigens, eggs in the stool or in rectal biopsy fragments), and imaging techniques (i.e. ultrasound with liver and/or spleen enlargement and lymph node adenopathy, magnetic resonance showing spinal cord injury). To be considered as having acute schistosomiasis in the present study, the participants had to have one or more of the symptoms/signs described above, evidence of an infection (parasitological or serologic), and reported contact with contaminated waters of the swimming pool. All 50 individuals fulfilled the criteria for the case definition of acute *S. mansoni* schistosomiasis and were treated with PZQ (60 mg/kg for children and 50 mg/kg for adults). From 50 individuals, 19 presented eggs in the feces after K-K examination (2 slides for each 2 stool samples) performed between 3 and 4 months after the date of contact with contaminated water. In this present study, 15 acute sera samples with egg-positive results were pooled for immunoproteomic analysis and classified as the INF-AC group.

#### Healthy individuals sera

Fifty five healthy individuals (35 women/20 men, 21-70 years old) living in a non-endemic area in Belo Horizonte, state of Minas Gerais were selected as donors to be used as our negative control group of individuals (NEG). They were interviewed and had no medical history of previous schistosomiasis. Parasitological examination was performed by K-K and GS as described. Serological reactivity to SEA and SWAP was performed by IgG-ELISA in the Reference Center for Schistosomiasis as previously described [35]. Patients with eggs in the feces and reactive for both IgG-ELISA assays were removed from the healthy group. In this present study, 15 sera samples were pooled for immunoproteomic analysis and all 55 sera samples from NEG group were used for standardization of the IgG-ELISA.

### Immunoproteomics analysis

#### Preparation of protein extract

The preparation of SEE was performed as Ashton et al. (2001) with modifications [56]. BALB/c mice female were infected by the subcutaneous route with 100 *S. mansoni* cercariae of the LE strain. After 45 days, they were sacrificed; their livers were removed, homogenized and digested with trypsin for 3 h at 37°C. After incubation, the livers were sieved and the eggs were collected by sedimentation and cleaned by washing 6 times in Phosphate-Buffered Saline (PBS). Cleaned eggs were re-suspended in 1 mL of Tris-Buffered Saline (TBS) supplemented with protease inhibitor cocktail (Sigma-Aldrich) and 1% dithiothreitol (DTT). The suspension was sonicated using six 10 sec pulses on full power with 1 min on ice between each sonication. Sonicated suspension was centrifuged at 100,000 g for 60 min and the supernatant collected. Protein concentration of SEE was measured by the Bicinchoninic Acid Assay (BCA) (ThermoScientific) and the quality of the extract was verified by SDS-PAGE 12%. Next, acetone precipitation was performed and the pellet was solubilized in rehydration buffer (7 M Urea, 2 M Thiourea, 2% 3-3-Cholamidopropyl-dimethylammoniopropane-sulfonate (CHAPS), 0.002% bromophenol blue) and stored at −70°C until use.

#### Two-dimensional-polyacrylamide-gel-electrophoresis (2D-PAGE)

Sixty micrograms of protein extract was used for 2D-PAGE to be stained in gel and 45 micrograms extract for 2D-PAGE to be used for Western blot. SEE proteins solubilized in rehydration buffer were supplemented with 1% DTT and 0.8% ampholyte 3–10 buffer (Bio-Lyte, Bio-Rad) and submitted to first dimension. Samples were loaded onto 7 cm immobilized pH gradient (IPG) strips, 3–10 pH ranges (Immobiline DryStrip Gels, GE Healthcare) for isoelectric focusing (IEF). IEF was conducted in Ettan IPGphor 3 (GE Healthcare) at 20°C and 50 μA/strip under the following conditions: passive in-gel rehydration at 50 V for 12 hs and focalization at 500 V for 30 min, followed by 1,000 V for 30 min and 8,000 V for 3 hs. Focused proteins were reduced and then alkylated using 1% DTT and 4% iodoacetamide, respectively, in equilibration solution (6 M urea, 75 mM Tris-HCl pH 8.8, 30% glycerol, 2% SDS, 0.002% bromophenol blue) for 15 min each at room temperature (RT). For the second dimension, IPG strips and molecular weight standards were then placed on top of 12% SDS-PAGE gels and sealed with 1% agarose. Electrophoretic protein separation performed using Mini-Protean III (Bio-Rad) at 20 mA/gel, for approximately 6 h. Gels were fixed in 2% v/v orthophosphoric acid/30% v/v ethanol solution overnight, then washed 3 × 10 min with 2% v/v orthophosphoric acid. Gels were stained with 2% v/v orthophosphoric acid/18% v/v ethanol/15% w/v ammonium sulfate/0.002% w/v Colloidal Coomassie Blue G-250 (Sigma-Aldrich) solution for 48 h. Gels were destained in 20% v/v ethanol for 5 min. For each experiment, three 2D-PAGEs were performed in parallel, one for Western blotting with native extract, another for western blotting with deglycosylated extract and another for stain and spot excision for protein identification.

#### Two-dimensional western blotting (2D-WB)

Proteins in 2D-PAGE were electrophoretically transferred to PVDF membrane 0.2 μm (GE Healthcare) using a Mini-Trans-Blot (Bio-Rad) at 100 V (2–3 mA cm2) for 2 h at 4°C with transfer buffer (25 mM Tris-Base, 192 mM glycine, 20% methanol). Post-transfer, PVDF membranes were stained with Ponceau for 10 min and quickly washed in water. Membranes were then blocked with TBS (20 mM Tris-HCl, 500 mM NaCl, pH 7.5) containing 0.05% Tween-20 and 5% skim milk (TBS-T/5% milk) at 4°C for 16 h. Membranes were washed five times, 5 min/wash, in TBS-T. To alter glycan structures on the membranes, we used the method of Woodward et al. (1985) [57]. Membranes were incubated in 10 mM Sodium Metaperiodate (SMP) solution in 50 mM acetate buffer, pH 4.5, at RT for 1 h in the dark. Membranes were then washed with acetate buffer for 10 min, then incubated in 50 mM sodium borohydride in PBS for 30 min at RT. After five, 5-min washes with TBS-T, membranes were individually incubated in pooled human sera. The native membranes and SMP-treated membranes were incubated separately for 2 h with each pool of INF-CR, HT, PT, and NEG sera diluted 1:600 in TBS-T/3% milk and INF-AC sera diluted 1:1200 in TBS-T/3% sera. After five, 5-min washes with TBS-T, the membranes were incubated with anti-human IgG peroxidase conjugated antibody (A0170, Sigma), diluted 1:100,000 in TBS-T/3% milk at RT for 1 h. After five, 5-min washes with TBS-T and one 10-min wash in TBS, the immunoreactive proteins were developed using ECL Plus Western Blotting Detection System (GE Healthcare) and images captured using chemiluminescence detection in ImageQuant LAS 4000 (GE Healthcare). The 2D-WB experiments were performed in duplicate.

#### In gel-digestion

The 2D-WB and its corresponding Coomassie stained 2D-PAGE were overlapped using software Photoshop (Adobe Systems Incorporated) and spots identified in duplicate experiments were selected for identification. The antigenic protein spots were manually and individually excised from the corresponding 2D-PAGE for mass spectrometry (MS) identification. Selected spots were destained in 40% ethanol/7% acetic acid at 37°C until clear. Gel pieces were then washed in ultrapure water and reduced in 50 mM DTT at 65°C for 30 min and then alkylated in 100 mM iodoacetamide, at RT for 1 h. Gel pieces were then washed in 20 mM ammonium bicarbonate (AB)/50% acetonitrile (ACN) for 3 × 20 min each and fully dried using Speed Vac Concentrator Plus (Eppendorf). Gel slices were rehydrated in 15 μL of the digestion buffer containing 0.01 μg/μL of Sequencing Grade Modified Trypsin (Promega) in 20 mM AB for 20 min. Excess trypsin was removed and additional 40 μL of 20 mM AB added. Trypsinolysis was performed for 48 h at 37 °C. Then digestion supernatants transferred to clean tubes and 50 μL of 0.1% trifluoroacetic acid (TFA)/50% ACN were added to gel slices for 30 min. The supernatants from both tubes containing the tryptic peptides were pooled, dried by speed vacuum, and ressupended in 10 μL of 0.1% TFA. The peptides were desalted in reverse phase micro-columns Zip Tip C18 (Millipore), according to manufacture instructions. Peptides were dried again and re-suspended in 20 μL of 0.1% TFA for liquid chromatography–mass spectrometry (LC-MS) analysis.

#### Protein identification by mass spectrometry

Digestion products were analyzed by liquid chromatography–mass spectrometry (LC-MS) on a Q-Exactive hybrid quadrupole-orbitrap mass spectrometer (Thermo Scientific). Four microliters of peptide samples were injected into a nano UHPLC instrument (Dionex UltiMate 3000, Thermo Scientific) through a trapping system (Acclaim PepMap100, 100 um × 2 cm, C18, 5 um, 100 A, Thermo Scientific) for 3 min using as solvent 98% water / 2% ACN with 1% TFA and subsequently directed into a capillary column (Acclaim PepMap100, 75 um × 25 cm, C18, 3 um, 100 A, Thermo Scientific). Reverse-phase separation of peptides was performed at 40°C in a gradient of solvent A (water, 0.1% formic acid) and B (80% ACN / 20% water, 0.1% formic acid), at a flow rate of 300 nL/min. Peptides were sequentially eluted over a gradient spanning from 3.2 % to 12% ACN over 2 min and from 12 % to 44 % ACN over an additional 15 min. Peptide ions were detected using positive mode through data dependent analysis. Resolution for precursor ions (MS1) was set to 70,000 (FWHM at 200 *m/z*) with an automatic gain control target of 3e^6^, maximum injection time of 100ms, scanning over 300-2000 *m/z*. The Top12 most intense precursor ions of each MS1 mass spectra were individually isolated with a 2.0 Th window for activation via higher-energy collisional dissociation (HCD) with normalized energy of 30 V. Only peptides exhibiting charge states of +2, +3, +4 and +5 were selected. Automatic gain control target was set to 5e^5^ (minimum accumulation of 3.3e^3^) with maximum injection time of 150 ms. Dynamic exclusion of 40 sec was active.

Spectral data were submitted to Proteome Discoverer v.1.4 (Thermo) for database search using SEQUEST HT against 10.779 *S. mansoni* protein sequences (5.136.273 residues). Search parameters included cysteine carbamidomethylation as a fixed modification, methionine oxidation and protein N-terminal acetylation as variable modifications, up to one trypsin missed cleavage site, error tolerance of 10 ppm for precursor and 0.1 Da for product ions. A quality filter was applied to keep False Discovery Rate (FDR) < 0.1. The average area for the 3 most intense peptides was used to infer protein abundance. This was particularly important when more than one protein identity was assigned to the same gel spot.

### Production of recombinant protein

#### Cloning, expression and purification of major egg antigen

The recombinant *S. mansoni* major egg antigen (rMEA) was produced by Gateway cloning technology (Invitrogen). First, the coding region from the gene of interest was obtained by PCR amplification using cDNA from *S. mansoni* eggs as template (BEI Resources, Catalog no. NR-49421, U.S. Government property). Based on the nucleotide sequence (Smp_049250.1, GeneDB) the primers were designed: forward (5’-ATGTCTGGTGGGAAACAACATAACGCA-3’) and reverse (5’-CTAGTGAGTAATCGCATGTTGCTTCTCCAATG-3’). PCR amplification included DNA polymerase buffer 20 μL, 10 mM dNTP mixture 5 μL, 100 μM forward primer 1.5 μL, 100 μM reverse primer 1.5 μL, 50 ng cDNA, Q5 High-Fidelity DNA polymerase 1 μL (New England Biolabs), RNase-free dH_2_O to 100 μL final volume. Secondly, the purified PCR product was used as template for the following PCR amplification which involved DNA fragments flanked by attB sites. The primers were designed with sites attB incorporated and underlined in fusion with N-terminal histidine tag following the Gateway manufacturer’s instructions [58]: forward (5’-<u>GGGGACAAGTTTGTACAAAAAAGCAGGCTTCGAAGGAGATAGA</u>ATGTCTGGTGGGAAACAACATAACGC-3’) and reverse (5’-<u>GGGGACCACTTTGTACAAGAAAGCTGGGTC</u>CTAGTGAGTAATCGCATGTTGC-3’). The PCR product with flanked attB sites were inserted into pDONR221 vector by BP recombination reaction at 25°C for 16 h yielding the entry clone. Next, the entry clone containing the gene of interest flanked by the attL sites was integrated into the destination vector pEXP1-DEST by LR recombination reaction at 25°C for 16 h, producing the final expression clone. After each recombination reaction, the clones were transformed into subcloning competent DH5α *Escherichia coli* (New England Biolabs) by heat shock. The positive colonies were selected by PCR and grown in Luria Bertani broth supplemented with 100 μg/mL ampicillin (LB-Amp). The vector constructions were purified by QIAprep Spin Miniprep Kit (Qiagen) and the DNA sequencing verified using the M13 primers for entry clone and T7 terminator primers for expression clone primers (Eurofins Genomics). The expression clone was transformed into competent BL21 (DE3) *E. coli* (Novagen) and grown in LB-Amp at 37°C for 12 h. The culture was diluted 100-fold in LB-Amp until achieving an absorbance of 0.6 in 600 nm. Protein synthesis was induced by addition of 1 mM isopropyl β-D-thiogalactoside (IPTG) at 37°C for 4 h. Cells were then harvested by centrifugation and re-suspended in 40 mL of lysis buffer (50 mM Tris, 0.5 M NaCl, 0.2 mM EDTA, 3% sucrose, 1% Triton-X and 10 mM imidazole). Subsequently, the cells were submitted to three 30 s-cycles of sonication and centrifuged at 5400 g for 20 min. The protein was purified by affinity chromatography on Ni-NTA column (HisPur Ni-NTA Spin Columns, Thermo) under native conditions (imidazole: binding 10 mM, washing 3 × 25 mM and 1 × 100 mM, elution 500 mM). The purification of rMEA was verified by SDS-PAGE and western blotting using an anti histidine tag. Fractions containing rMEA were dialyzed against PBS pH 7.0 and concentrated using 30 kDa centrifugal tubes (Millipore). The recombinant proteins were quantified using BCA method and send to LC-MS to analysis by Shotgun.

#### Western blotting analyzes

Two, three and seven micrograms of rMEA were transferred onto 0.2 μm PVDF membrane strips after electrophoresis. The strips were blocked at 4°C for 16 h in TBS-T/3% skim milk and subsequently incubated separately with INF-CR and NEG sera diluted 1:800 at RT for 2 h. After five 5-min washes with TBS-T, the strips were incubated with anti-human IgG peroxidase conjugated antibody diluted 1:80,000 in TBS-T/3% milk at RT for 1 h. The strips were washed and revealed as 2D-WB.

### Application of rMEA in immunodiagnostic assay

Recombinant antigen rMEA was evaluated for the ability to diagnose *S. mansoni* infection by antigen-specific IgG ELISA (rMEA-IgG-ELISA). Optimization of the protocol and dilution of reagents were determined by titration. After standardization, the assay was performed in flat bottom plates (Maxisorp NUNC) sensitized with 100 μL/well of carbonate bicarbonate buffer 0.05 M, pH 9.6, containing rMEA at a concentration of 1μg/mL. The plates were incubated at 4°C for 16 h. Then plates were washed six times in PBS with 0.05% Tween 20 (PBS-T) the plates were blocked with PBS-T containing 2.5% skim milk at 37°C for 2 h. Then individual sera from INF-CR (n = 93) and NEG (n = 65) groups were diluted 1:100 in PBS-T and plated at 100 μL/well. Plates were incubated at RT for 2 h and washed six times in PBS-T. Peroxidase conjugated Anti-human IgG antibody was then added to wells at a dilution of 1: 60,000 in PBS-T at RT for 1 h. After 6 washes, plates were developed using 3, 3′, 5, 5′-tetramethylbenzidine (TMB, Sigma). The reaction was stopped with 50 μL of sulfuric acid and the optical density (OD) determined by an automatic ELISA reader (Multiskan, Thermo Scientific), using a filter of 450nm.

### Statistical analysis

Analyses were performed using Open Epi, version 3.03 and GraphPad Prism, version 5.0. In order to evaluate the performance of rMEA-IgG-ELISA, a Reference Standard was established, which included all positive results (visible eggs) from any of the parasitological methods used (K-K and SG). Normal distribution of the data was verified by the Shapiro-Wilk test. To compare the means for no normal distribution, the Mann-Whitney test was used with a p-value ≤ 0.05 considered significant. Receptor Operating Characteristic curves (ROC curves) were used to calculate area under curve (AUC), sensitivity, specificity and the cutoff points between infected (INF-CR) and healthy groups (NEG). The AUC indicates the probability of accurately identifying true positives, where one could distinguish between non-informative (AUC=0.5), less accurate (0.5<AUC≤ 0.7), moderately accurate (0.7<AUC≤ 0.9), highly accurate (0.9<AUC<1) and perfect tests (AUC =1) [59]. Positive predictive values (PPV), Negative Predictive Values (NPV) and overall accuracy (ACC) was determined by the following formula: PPV = number of true positives/(number of true positives + number of false positives); NPV = number of true negatives/(number of true negatives + number of false negatives) and ACC = (number of true positives + number of true negatives)/ (number of true positives + true negatives + number of false positives + number of false negatives).

The McNemar’s chi-square test was used to analyze categorical variables. To evaluate the degree of concordance between the different methods, the kappa index (κ) followed the categorization for Landis and Koch (1972): <0 poor, 0.00-0.20 slight, 0.21-0.40 fair, 0.41-0.60 moderate, 0.61-0.80 substantial and 0.81-1.00 almost perfect. The relationship between the intensity of infection (epg) determined by parasitological tests and the IgG-ELISA (OD) was examined by the Spearman correlation test.

## Results

### Antigens recognized in 2D analysis by pooled human sera in native and SMP-treated SEE

The 2D-PAGE provided good resolution of spots in pH range with minimal streaking. In order to identify the antigens recognized by antibodies in pooled sera, a corresponding 2D-PAGE was performed in parallel so that WB (native and SMP-oxidized) could be performed to exclude any variation that might arise from the use of different antigen preparations (Fig 1).

**Fig 1.**
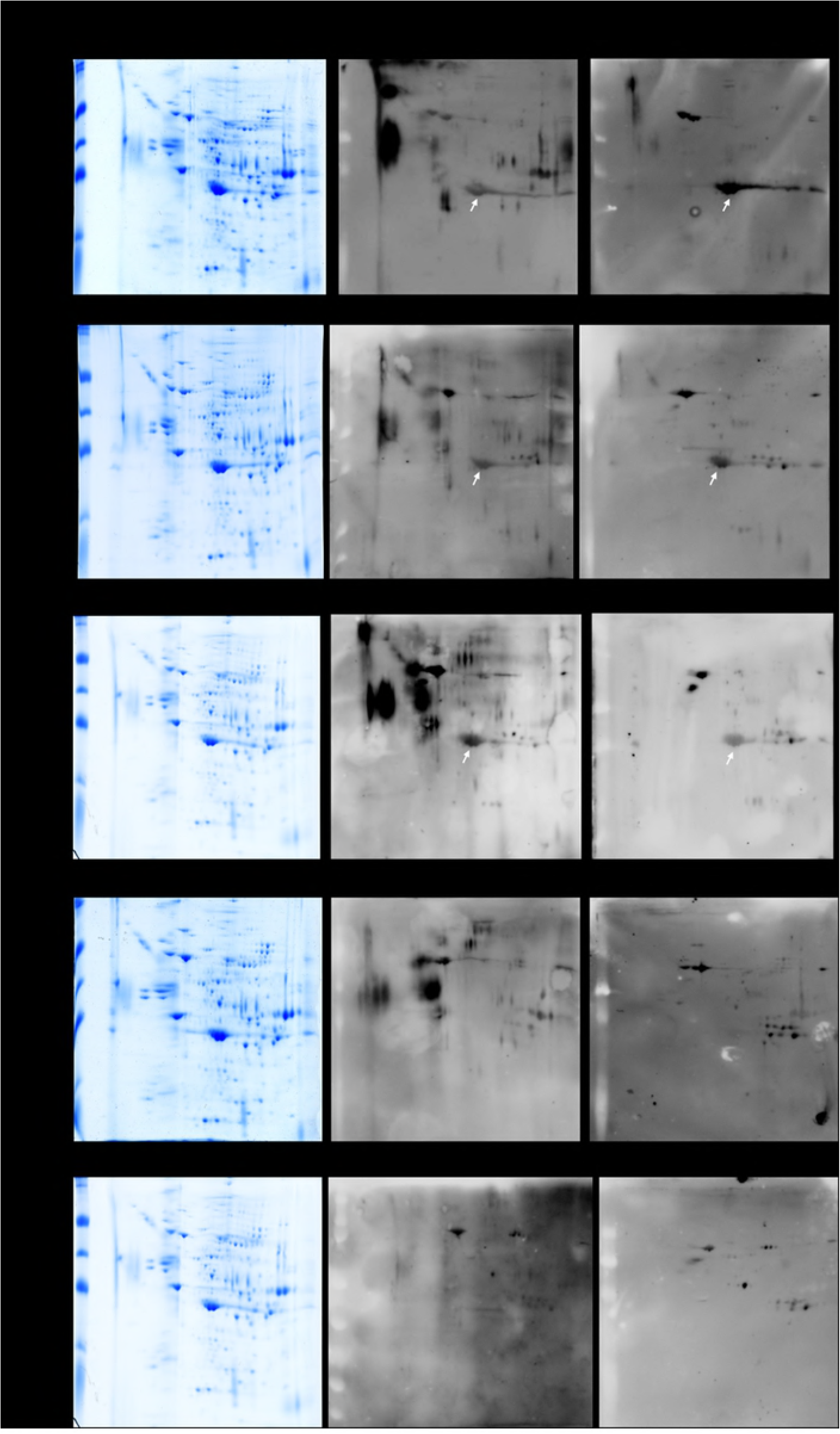
Two-dimensional analysis using *Schistosoma mansoni* egg extract and pooled sera from infected and non-infected individuals. A) 2D-PAGE of native SEE in 7 cm, pH 3–10 strip and stained by Coomassie G-250. B) Corresponding 2D-WB with pooled human sera and C) 2D-WB post membrane treatment with 10 mM of SMP. Sera from INF-CR, PT-CR, NEG and HT was added at 1:600 dilution. The INF-AC was added at 1:1200 dilution. The anti-human IgG peroxidase conjugate was added at 1:100,000. The white narrows indicate spot 5 corresponding to Major Egg Antigen.

In native 2D-WB, 23 immunoreactive spots were recognized by the pooled infected sera from *S. mansoni.* No difference in recognition was seen between INF-AC and INF-CR groups. From 23 spots, 22 spots were simultaneously recognized by HT and 10 were recognized by NEG group. One single spot, number 5 (indicated by white arrow on Fig 1), was exclusively recognized by infected patients (INF-AC and INF-CR) and was not recognized by the HT and NEG groups. Spot 5 was detected by antibodies in the pooled PT-CR group. The immunoblot and homologously stained gel were aligned and the 23 spots matched and excised for LC/MS analysis (Fig 2). The identification of spots is presented in Table 2.

**Fig 2.**
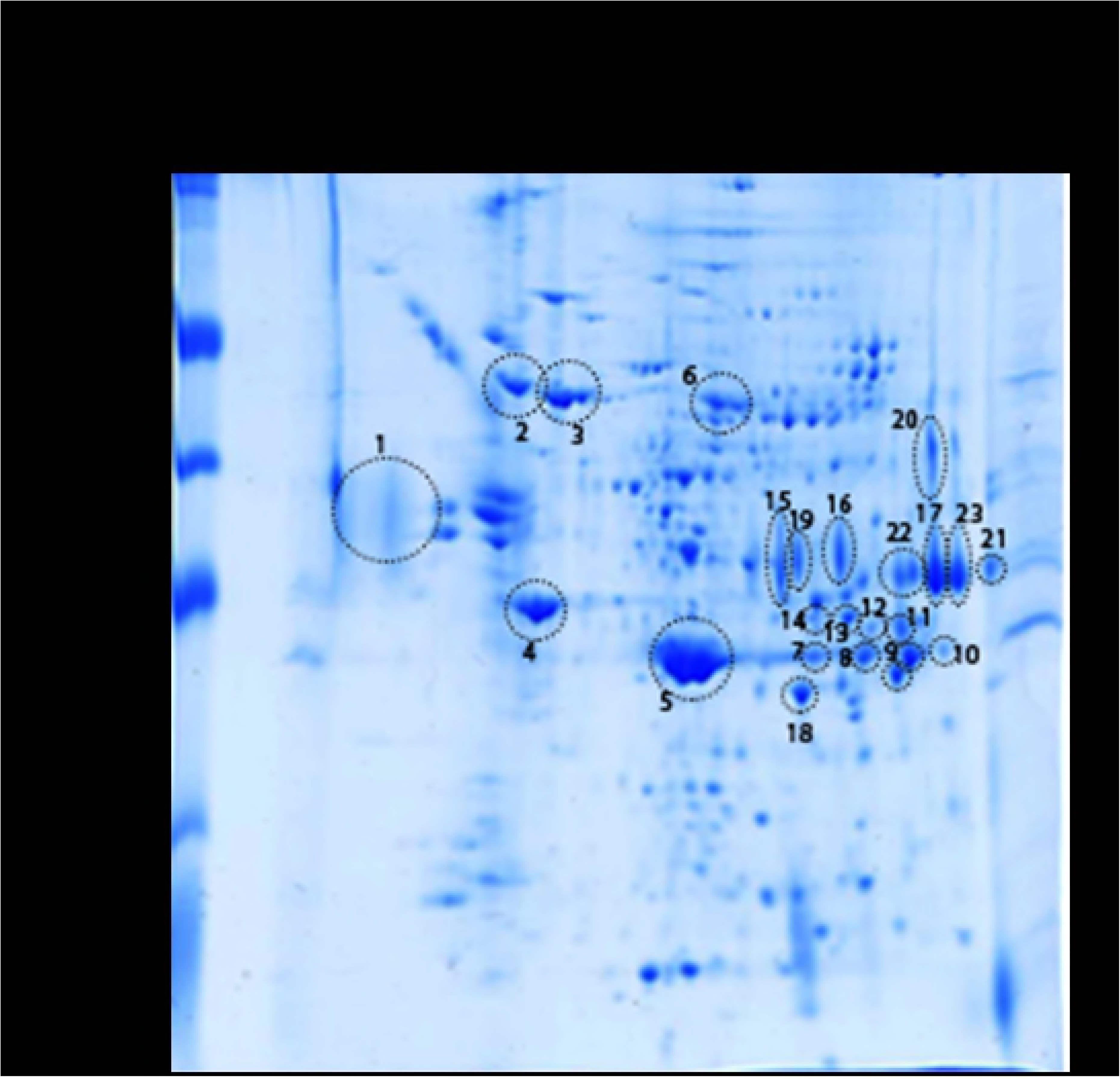
Coomassie blue-stained 2D-PAGE showing spots matched to the 2D-WB. Proteins from SEE were separated in 7 cm, pH 3–10 strip and stained by Coomassie G-250. Immunoreactive spots from infected *S. mansoni* sera were numbered (n = 23) and were excised and submitted for mass spectrometry identification.

**Table 2.** Identities of proteins recognized by Schisosoma-infected serum (acute and chronic) in soluble egg extracts

| Spots | Accession | Description | Unique peptides | Coverage (%) | MW (kDa) | pI | Score | Biological function |
| --- | --- | --- | --- | --- | --- | --- | --- | --- |
| 1 | Smp_170410.1 | NADH: ubiquinone oxidoreductase complex I | 7 | 25.83 | 29,2 | 4.7 | 37.83 | EM |
| 2 | Smp_049550.1 | 78-kDa glucose regulated protein | 17 | 32.87 | 71,2 | 5.22 | 132.4 | EM |
| 3 | Smp_106930.1 | Heat shock 70 kDa protein homolog | 24 | 48.04 | 69,8 | 5.58 | 229.93 | CH |
| 4 | Smp_183710.1 | Actin | 4 | 44.15 | 41,7 | 5.48 | 151.51 | ST |
| 5 | **Smp_049250.1** | **Major egg antigen** | 14 | 48.20 | 37,0 | 6.96 | 364.77 | CH |
| 6 | Smp_005880.1 | Phosphoenolpyruvate carboxykinase | 27 | 50.96 | 70,3 | 6.90 | 114.13 | EM |
| 7 | Smp_056970.1 | Glyceraldehyde-3-phosphate dehydrogenase | 3 | 10.36 | 36,4 | 8.05 | 7.89 | EM |
| 8 | Smp_056970.1 | Glyceraldehyde-3-phosphate dehydrogenase | 9 | 30.18 | 36,4 | 8.05 | 22.18 | EM |
| 9 | Smp_056970.1 | Glyceraldehyde-3-phosphate dehydrogenase | 12 | 37.87 | 36,4 | 8.05 | 33.84 | EM |
| 10 | No significant score* | --- | --- | --- | --- | --- | --- | --- |
| 11 | Smp_150240.1 | Secretory glycoprotein k5 | 3 | 3.76 | 31,1 | 8.19 | 6.67 | ST |
| 12 | Smp_079920.1 | Pyruvate dehydrogenase | 2 | 5.34 | 44,0 | 8.46 | 1.73 | EM |
| 13 | Smp_042160.1 | Fructose-bisphosphate aldolase | 8 | 23.15 | 35,4 | 8.06 | 27.89 | EM |
| 14 | Smp_042160.1 | Fructose-bisphosphate aldolase | 1 | 2.78 | 35,4 | 8.06 | 2.23 | EM |
| 15 | Smp_179250.1 | Alpha galactosidase: alpha n | 15 | 30.84 | 108,4 | 7.37 | 105.84 | EM |
| 16 | Smp_179250.1 | Alpha galactosidase:<br>alpha n | 2 | 22.93 | 47,7 | 7.53 | 42.76 | EM |
| 17 | Smp_150240.1 | Secretory<br>glycoprotein k5 | 3 | 9.40 | 31,1 | 8.19 | 25.98 | ST |
| 18 | Smp_035270.1 | Cytosolic malate<br>dehydrogenase | 12 | 36.27 | 31,3 | 8.48 | 55.54 | EM |
| 19 | No significant<br>score* | --- | --- | --- | --- | --- | --- | --- |
| 20 | No significant<br>score* | --- | --- | --- | --- | --- | --- | --- |
| 21 | No significant<br>score* | --- | --- | --- | --- | --- | --- | --- |
| 22 | Smp_150240.1 | Secretory<br>glycoprotein k5 | 3 | 15.41 | 31,1 | 8.19 | 41.09 | ST |
| 23 | Smp_150240.1 | Secretory<br>glycoprotein k5 | 2 | 9.02 | 31,1 | 8.19 | 23.59 | ST |

(\*): No significant score based on SEQUEST HT search engine output statistic. EM:

The 23 spots were resolved into 12 proteins. LC/MS analysis revealed instances in which different spots were derived from the same protein: for example, spots 11, 17, 22, and 23 are all secretory glycoprotein k5. It was observed that, in some cases, there was no direct correlation between the amount of protein in the SEE protein extract and its antigenicity level. Although most of the immunoreactive spots recognized by infected serum were visible in the corresponding 2D-PAGE, there were highly immunoreactive spots that were barely visible in stained gels (e.g., spot 1). Spots 10, 19, 20 and 21 were not identified due to low abundance. The most identified proteins were related to housekeeping proteins. These include structural/muscle proteins, enzymes (mostly components of the glycolytic pathway) and chaperone proteins.

To evaluate the presence of glycosylated epitopes on the 23 immunoreactive spots, 2D-WB were performed using SMP-treated membranes and then compared to the native one (Table 3). After oxidation, only 12/23 spots maintained immunoreactivity, indicating they potentially have protein epitopes. From these 12 spots, 11 spots cross-reacted with the HT and 10 spots cross-reacted with the NEG group. Spot number 5 was uniquely recognized by schistosome-infected groups (INF-CR and INF-AC) and was not recognized from uninfected groups (HT and NEG). Furthermore, there was an observed decrease in reaction intensity of spot 5 in PT-CR compared to the corresponding INF-CR group at baseline in the SMP experiment.

**Table 3.**
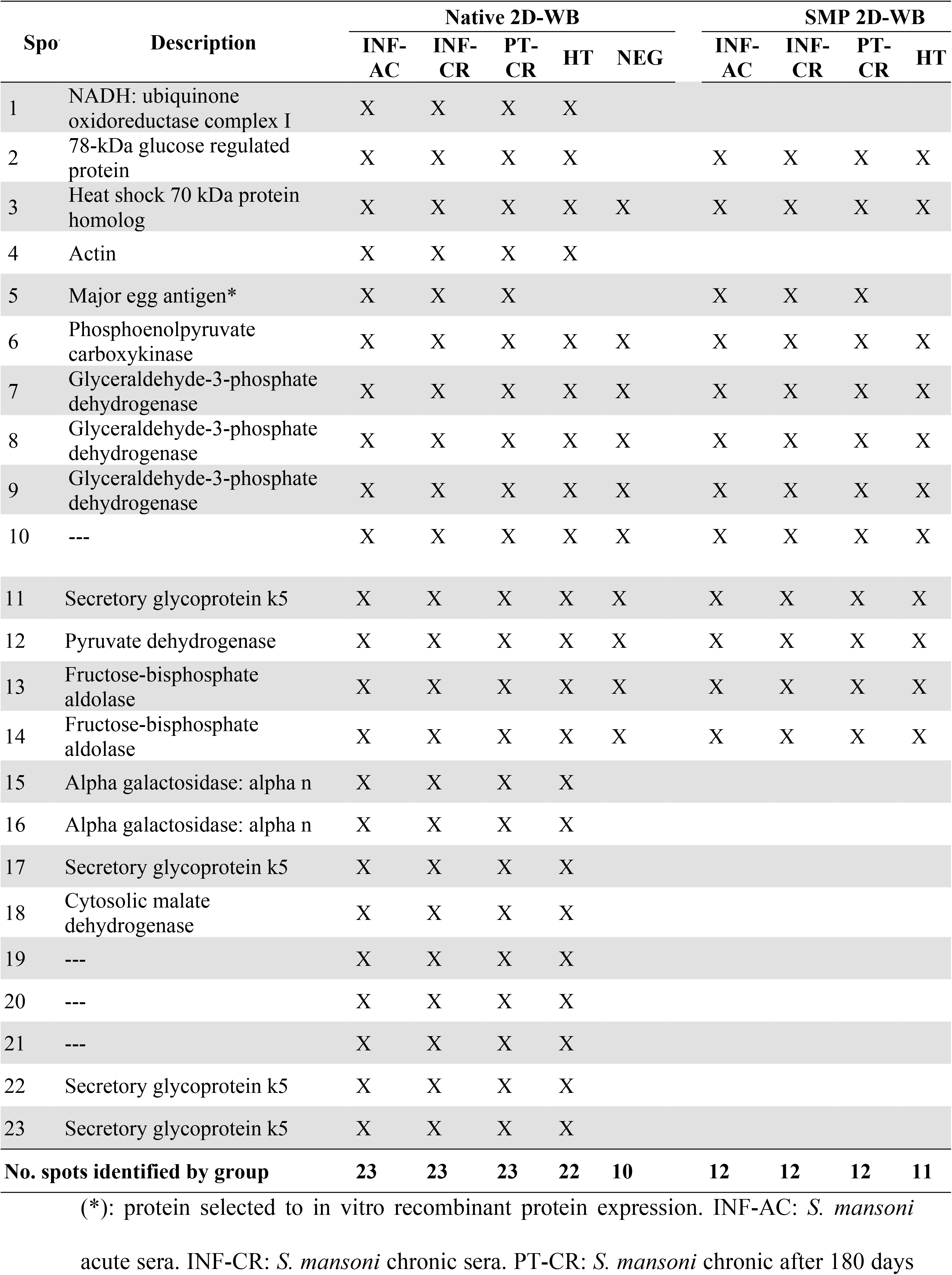
Comparative spot recognition by schisosome-infected and uninfected serum in soluble egg extract before and after sodium metaperiodate oxidation.

Spot 5, approximately 40 kDa and pI 7.0, was identified as Major Egg Antigen and chosen for further evaluation in immunodiagnostics assays. Selection was based on: 1) single identification in infected *S. mansoni* group (INF-AC and INF-CR), 2) absence of cross-reaction in *S. mansoni* uninfected groups (HT and NEG), 3) recognition after SMP treatment (potential presence of immunogenic peptides and feasibility for bacterial production) and 4) decrease of reactivity intensity in PT-CR group.

### Expression and purification of rMEA

The rMEA was expressed by IPTG induction in *E. coli*. The size from recombinant construction was predicted in Expasy Software including the histidine tag (https://www.expasy.org/proteomics/protein_structure) corresponding to 43 kDa. As shown in Fig 3, the purified protein was present in the gel and the corresponding western blotted anti-histidine tag. To validate the recombinant proteins, the purified material was sent for MS analysis by Shotgun. The results showed 98.6% of abundance was related to native MEA (Smp_049250.1), confirming the identity.

**Fig 3.**
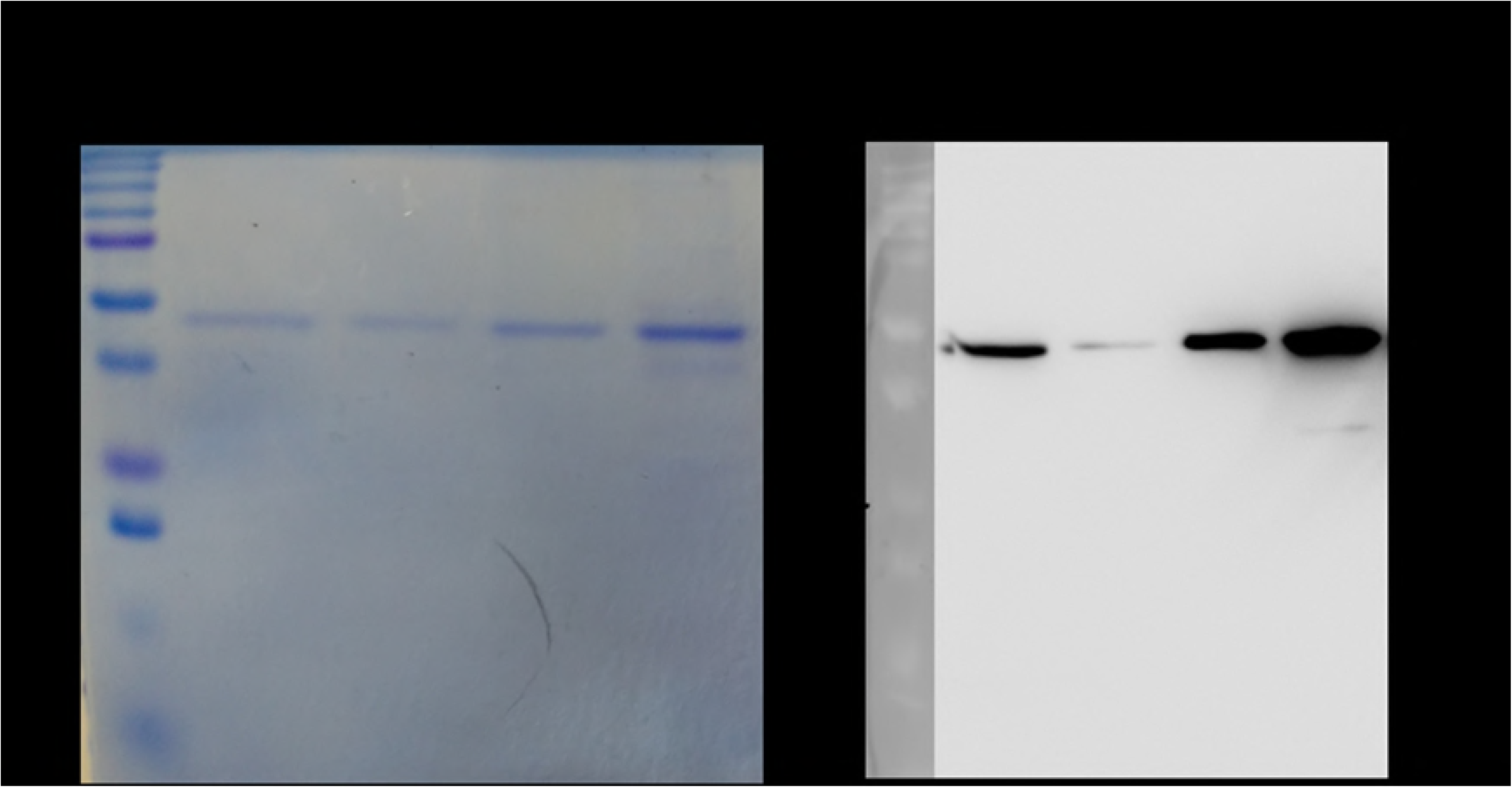
Coomassie blue-stained SDS-PAGE and western blotting anti-his tag with the purified rMEA. Purified rMEA was run in 12% gel (A). Replicate gel was transferred to PVDF membrane and probed with mouse anti-histidine tag followed of anti-IgG conjugated with peroxidase.

We evaluated the antigenicity from rMEA using serum from *S. mansoni* infected individuals from endemic areas and non-infected healthy individuals (NEG). The recombinant protein maintained the recognition pattern in the 2D-WB experiments, confirming their correct identification and the maintenance of antigenic epitopes in the *in vitro* expression (Fig 4). After all confirmation, rMEA was followed to diagnosis evaluation by IgG-ELISA.

**Fig 4.**
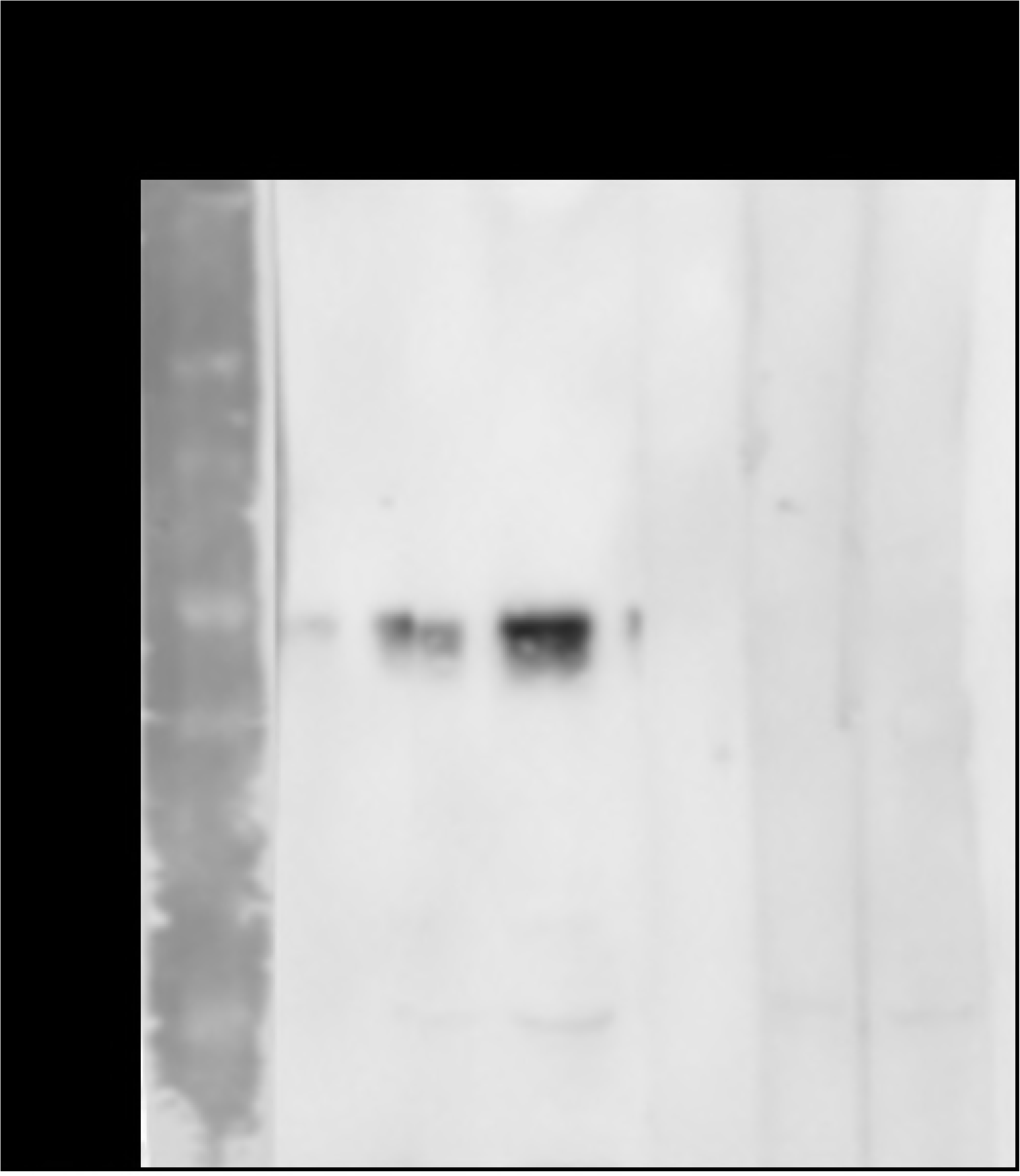
Western Blotting anti-rMEA using serum from infected *S. mansoni* individuals and non-infected individuals. Three different concentrations of rMEA was run in 12% gel. The strips were probed with serum from INF-CR and NEG group at 1:800 dilution. The anti-human IgG peroxidase conjugated was added at 1:80,000.

### Assessment of rMEA in diagnosis of schistosomiasis

The ROC curve analysis for rMEA-IgG-ELISA was carried out to estimate the cutoff and performance indices (sensitivity, specificity, PPV, NPV and AUC), using the NEG and INF-CR groups as reference groups. The AUC demonstrated a high power of discrimination between the groups (AUC = 0.95). The cut off was 0.232 and selected based on the best overall accuracy (ACC = 87.8%). The sensitivity was 87.10% and specificity was 89.09% with PPV and NPV of 93.1% and 80.33% respectively. Significant IgG reactivity against rMEA was observed in *S. mansoni* infected individuals (INF-CR) in comparison with negative healthy donors (NEG) and those negative from endemic areas (NEG-END) (Fig 5).

**Fig 5.**
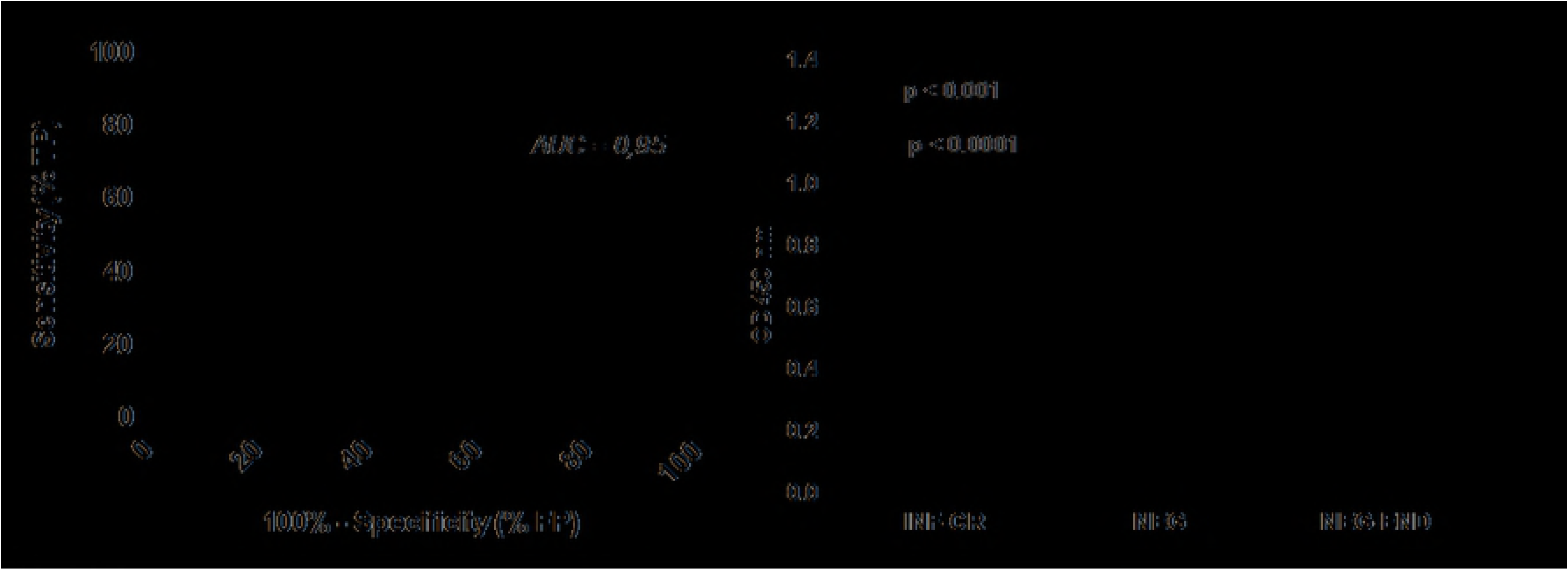
ROC analysis and human IgG-specific response against rMEA. ROC was carried out with 93 positive sera samples from low endemic areas (INF-CR) and 55 negative sera samples from non-endemic areas (NEG). Additionally, 80 samples from individuals with negative stool examination from endemic areas (NEG-END) were evaluated. Significant differences between groups are indicated in the graphic (Mann Whitney test, CI: 95%). Dashed lines represent the cutoff in the absorbance level, which determines specificity and sensitivity.

From 93 egg-positive individuals, the immunoassay was able to identify 81 infections of which 75 had low-intensity infection (<100 epg). From 64 individuals harboring extremely low-intensity infections (≤ 10 epg), rMEA-IgG-ELISA identified 56 infections of which 27 intensity of infection at 1 epg (Table 4). The parasite load from 12 individuals varied from 1 to 99 epg. There was not a significant positive correlation between the IgG levels (OD) and the parasitological load (epg) by Spearman rank test (r = 0.024, p = 0.8167). From 80 stool negative individuals from the endemic area, rMEA-IgG-ELISA identified 41 as positive.

**Table 4.** Sensitivity of rMEA-IgG-ELISA for the detection of schistosomiasis considering the parasite load, as defined by egg counts of 2 grams of feces (24 Kato-Katz slides and 2 Saline Gradient).

| Low-intensity (1-99 epg) |  |  |  | Moderate (100-399 epg) |
| --- | --- | --- | --- | --- |
| $\leq 10$ | 11-25 | 26-50 | 51-99 | $\geq 100$ |
| % Sensitivity | % Sensitivity | % Sensitivity | % Sensitivity | % Sensitivity |
| 87.5 (56/64) | 83.3 (10/12) | 66.7 (2/3) | 87.5 (7/8) | 100.0 (6/6) |

The agreement between rMEA-IgG-ELISA and the reference method determined by 24 slides of K-K and 2 procedures of SG showed substantial concordance (κ = 0.75). When the current adopted K-K (2 slides) was compared, it demonstrated a fair concordance (κ = 0.32) underdiagnosing 57 true cases. The positivity rate from rMEA-IgG-ELISA (58.8%) and the reference method (62.8%) showed no significant difference (χ^2^, p = 0.24) (Table 5).

**Table 5.** Comparison of positive and negative results determined by rMEA-IgG-ELISA and parasitological analysis (saline gradient or Kato-Katz, 2 or 24 slides).

| rMEA-IgG-ELISA | Parasitological (2000 mg feces)* |  |  |  |  |  | Parasitological (83.4 mg feces)** |  |  |  |  |  |
| --- | --- | --- | --- | --- | --- | --- | --- | --- | --- | --- | --- | --- |
|  | Positive |  | Negative |  | Total |  | Positive |  | Negative |  | Total |  |
|  | N | % | n | % | N | % | n | % | n | % | n | % |
| Positive | 81 | 87.1 | 6 | 10.9 | 87 | 58.8 | 36 | 38.7 | 0 | 0.0 | 36 | 24.3 |
| Negative | 12 | 12.9 | 49 | 89.1 | 61 | 41.2 | 57 | 61.3 | 55 | 100.0 | 112 | 75.7 |
| Total | 93 | 62.8 | 55 | 37.2 | 148 | 100 | 93 | 62.8 | 55 | 37.2 | 148 | 100 |
| Sensitivity: 87.10% (78.55% to 93.15%) |  |  |  |  |  | Sensitivity: 38.71% (28.78% to 49.38%) |  |  |  |  |  |  |
| Specificity: 89.09% (77.75% to 95.89%) |  |  |  |  |  | Specificity: 100.0% (93.51% to 100.0%) |  |  |  |  |  |  |
| Kappa = 0.75 |  |  |  |  |  | Kappa = 0.32 |  |  |  |  |  |  |
| p = 0.24 |  |  |  |  |  | p = 0 |  |  |  |  |  |  |

\* Analysis by 24 slides of K-K and 2 procedure of SG
\*\* Analysis by 2 slides of K-K
Significant differences expressed by McNemar's chi-square test.

## Discussion

Advances in development of new schistosomiasis diagnostic methods are necessary for low prevalence/low-intensity infections. In the majority of Brazilian endemic areas, transmission is maintained by individuals harboring low level infections that are undiagnosed by analysis of 2 slides of K-K in a single stool sample [6]. It has been suggested that a diagnostic test with high specificity and sensitivity will be needed for prevention, control and elimination of this disease [7, 12]. Some diagnostic methods have already shown their ability to detect positive individuals missed by parasitological techniques, due to low or absence of egg excretion [13, 14, 19]. Immunoassays based on antigen or antibody detection have been the most studied, due to high sensitivity and ease-of-translation to a format applicable to field settings [60].

The POC-CCA^®^ is the most recently evaluated test to be part of WHO guidelines. This immunochromatographic RDT has shown good performance for mapping areas and assessing cure after treatment in Africa [61]. However, the studies conducted in Brazil were controversial. The issues about adopting POC-CCA^®^ in Brazil are related to inadequate interpretation of trace as positive following the manufacturer’s recommendation and the low sensitivity to detect low-intensity infections [11, 12, 20, 27-33]. The authors emphasized differences regarding the predominance of extreme low-intensity infections (1-25 epg) found in the Brazilian areas [10-12].

Antibody-based immunodiagnostics have greater sensitivity than parasitological methods in low-endemic areas [14, 19, 34-39]. Serum immunoreactivity to schistosome antigens allows detection of infections with loads as low as 1 epg. Even though the antibody-detection methods are currently not the first choice for endemic areas, due to their inability to discriminate current infection from previous exposure, as well as the inability to monitor treatment effectiveness, they have been accepted as a complementary tool during epidemiological survey [13, 19, 34, 62]. One of the difficulties in developing antibody-detection methods is the choice of antigen. Crude antigens, such as SEA and SWAP, are not ideal in terms of sensitivity and specificity. To overcome this, the search for new antigens and subsequent production by recombinant strategies has been proposed as an alternative to improve antibody detection. In this study, we aimed to identify parasite markers for development of highly sensitive and specific new immunological tests. By using serum from individuals from a low-endemic area in Brazil in 2D-WB, we were able to screen antibody targets directed to egg extracts and identify an antigen, which shows potential for diagnosis of low-intensity schistosomiasis infections.

Our work was the first serological-proteomic study conducted with egg extracts from *S. mansoni* and human samples. Further, we included diversified sets of sera allowing for a more rational search for a highly specific diagnostic marker. Through the immunoproteomic approach, we identified 12 different immunogenic proteins from egg extracts. Other *Schistosoma spp.* serological-proteomic studies using human samples have been conducted. Mutapi et al. (2005) used serum from infected individuals with *S. hematobium* to screen adult worm antigens in two-dimensional electrophoresis (2-DE) to identify suitable antigens for diagnostic purposes. Twenty six immunoreactive protein spots were identified and investigated [51]. The unique study related to *S. mansoni* and human samples involved searching for vaccines candidates using worm extract. Ludolf et al. (2014) identified 47 different immunoreactive proteins from worm antigens using sera from positive and negative endemic individuals. One of them, the eukaryotic translation elongation factor, uniquely reacted with naturally resistant residents from endemic areas and was considered a potential vaccine candidate [52].

Our results showed that 23 immunoreactive spots, resolved into 12 different proteins, were strongly recognized by pooled sera from schistosome-infected individuals. No differences were found between acute and chronic samples. Currently, differentiation between the two stages of infection is based on clinical and epidemiological data. Differentiating them by serological diagnosis could contribute to the establishment of adequate protocols for treatment of infected patients and detection of new foci or infection cases in tourists. However, this work did not identify proteins specific for different stages of infection. Some studies initially pointed out antigens, such as SmRP26 and KLH, with the potential to discriminate between the acute and chronic phase, however, there was no reproducibility in subsequent evaluations [63-65]. A more sophisticated study using protein microarrays evaluated the recognition of 92 proteins in sera from infected individuals (acute and chronic phase) and negatives from a non-endemic area. Fifty antigens were recognized by sera samples in the acute and chronic phase. From these, 4 antigens were differentially recognized between the acute phase and chronic phase and will be further evaluated in the standardization and validation of new differential methods for the diagnosis of different infection stages [66].

Differential recognition was not found between the infected group and post-treatment group. Antibodies remain present in serum following treatment of infected individuals, making it difficult to differentiate between current and previous infections [43]. The persistence of antibodies after treatment impairs post-treatment monitoring, which could be resolved by means of a differential diagnosis using an antigen specific for that phase. Mutapi et al. (2005), using a similar approach to this work, but with *S. haematobium* infections, identified 5 exclusively immunoreactive proteins in serological samples after treatment. The presence of new antigens at this stage was related to the release of these antigens after parasite death and exposure to the host’s immune system [51].

To narrow down our search, we added 2D-WB analysis and further treated some of the membranes with SMP to identify spots that were likely proteins. Periodate oxidation is employed to alter glycan structures and therefore eliminate their ability to be detected by anti-glycan antibodies on glycoproteins [57]. This is important as polyparasitism is common in endemic areas and glycans are the most shared and most immunogenic fractions among helminth species [54, 56, 67]. From the initial 23 spots recognized in native extract by Schistosoma infected patients, 22 were shared with helminth-infected patients. After oxidation with SMP, the number of spots for Schistosoma-infected patients decreased to 12 and the helminth-infected patients decreased to 11 spots. No difference was noticed in recognition by negative sera. Although not all carbohydrates are sensitive to periodate treatment [57], these data suggest the influence of carbohydrate moieties on the immunogenicity of glycoproteins and also their participation in cross-reactions with other parasites. Alarcón de Noya et al. (2000) demonstrated that after oxidation of egg extracts, the specificity of the IgG-ELISA test increased in the detection of *S. mansoni*-infected individuals [44].

Among the proteins identified in this study, the Major Egg Antigen was selected to be produced in a bacteria model and the recombinant evaluated in the immunodiagnostic for schistosomiasis. It was the unique antigen that was recognized by Schistosoma infected patients, but was not recognized by negative individuals and those infected with other helminths in 2D-WB. Further, MEA maintained immunoreactivity after SMP treatment. MEA, also known as Smp-40, is one of the 40 most abundant proteins secreted by the eggs [67-69] and can be found in adult worms [70]. MEA is a chaperone and shares homology with the family of heat shock proteins. It is involved in the protection of miracidia from oxidative stress, denaturation, and aggregation of proteins [67]. MEA has been described as highly antigenic in infected humans [71]. The profile of cytokines obtained from peripheral blood mononuclear cells (PBMCs) from *S. mansoni* infected patients and stimulated with purified MEA was associated with reduced granuloma formation and an anti-pathological vaccine [72].

MEA was previously suggested as a diagnostic by Nene et al. (1986), since it was immuno-precipitated in human serum [73]. Ludolf et al. (2014) demonstrated the immunoreactivity of recombinant MEA antigen against samples from chronic individuals using western blotting [52]. In this study, rMEA was produced and recognized by sera from infected endemic individuals and was not recognized by sera from negative non-endemic individuals in WB analysis. Since this antigen proved to be promising in preliminary WB, we evaluated the performance of rMEA in the detection of IgG by ELISA.

Although antibody detection cannot differentiate between active and past infection, its main advantage is in the detection of individuals recently exposed (pre-patent phase) and those with low parasitic loads that are not detected using K-K (2 slides) [13, 14, 19, 74]. Among serological tests, ELISA is widely used for the diagnosis of schistosomiasis, due to its low relative cost and ability to run a lot of samples at the same time. In an outbreak in Southeast in Brazil, the IgG-ELISA was used to diagnose tourists from a non-endemic area recently exposed to an infected river considered an unknown focus of transmission. Kinkel et al. (2012) evaluated 8 serological assays and addressed to them the value for diagnosis of schistosomiasis in individuals from areas where the disease is not endemic and who are carrying light and/or recently acquired infections [42]. In endemic regions, residents are continuously exposed to parasite infection and parasite antigens. However, many have high titers of antibodies without being infected. Therefore IgG-ELISA can be useful as a screening tool. Pinheiro et al. (2012) carried out the screening of individuals from a low-endemic area in Brazil through detection of antibodies. Positive individuals in the serology were subject to an extensive parasitological evaluation (K-K 24 slides, Helmintex and SG) [14]. In the study by Da Frota et al. (2011), 85 egg-negative, but IgG-positive cases were evaluated a second time with additional samples and K-K slides and the positivity increased from 3.8% to 8.7% [19]. Espírito-Santo et al. (2015) adopted IgG and IgM-ELISA as a screening tool followed by analysis by a more specific method (PCR) in the detection of cases in a low-endemic area in Brazil. This algorithm showed good performance to determine the true prevalence compared to analysis by 2 K-K slides [13].

Recombinant antigen ELISAs have potential to increase the specificity of assays [74-77]. In our study, rMEA-IgG-ELISA showed sensitivity of 87.10% and specificity of 89.09% represented by AUC = 0.95. AUC is a measure of diagnostic precision in which the value of 1 indicates a perfect test and 0.5 is non-discriminant [59]. Our ROC curve analyses showed highly accurate power of discrimination indicating almost perfect test. Using a similar set of samples, Sarhan et al. (2014) showed AUC of 0.99 and 0.87 using SEA and SWAP antigens [78], and Grenfell et al. (2013) showed 0.94 using recombinant CCA [79]. Even crude extracts of worm antigens can be used for ELISA, such assays would require infrastructure to maintain the parasite cycle and the complexity of large-scale production and standardization. Furthermore, the low performance in regards to sensitivity and specificity from crude antigens remains a problem.

The rMEA-IgG-ELISA performed significantly better than the currently adopted K-K (2 slides) for detection of low-intensity infections. The K-K exhibited a sensitivity of 38.71% with 57 false negative cases, ranging from 1-10 epg. In this case, undiagnosed individuals would not receive treatment, possibly develop serious disease, and contribute to maintenance of transmission. On the other hand, rMEA-IgG-ELISA showed 87.5% (56/64) of sensitivity in the group of ≤ 10 epg, which is indicative of the majority of cases of schistosome infection in Brazil [10-12, 20, 27, 28]. Additional studies utilizing other *S.mansoni* recombinant proteins also showed better results than K-K performed by 2 slides in low-endemic areas. The recombinant CCA showed 100% sensitivity and 96% specificity by IgG detection in chronic individuals using magnetic microspheres [79]. The IgG-ELISA using recombinant 200-kDa tegumental protein demonstrated 90% sensitivity and 93.3% specificity with a strong correlation with egg burden in the same set of individuals [75]. El Aswad et al. (2011) showed sensitivity and specificity of 89.7% and 100%, respectively, using the recombinant calreticulin and cercarial transformation fluid in ELISA [80]. The rMEA-IgG-ELISA did determine that a number of negative residents from endemic areas were positive. This issue has been reported in other studies discussing single test immunoassay false positives and why any single assay may not be appropriate for epidemiological surveys [53, 80-82].

In the work presented here, we demonstrated that the immunoproteomic analysis was successful in selecting a good candidate for use in the diagnosis of schistosomiasis. The rMEA showed performance in ELISA superior to the current gold standard K-K since high levels of IgG were identified in individuals harboring 1-10 epg missed in primary parasitological analysis (K-K 2 slides). We believe the IgG-ELISA can be a useful tool to be combined with other techniques in low-endemic areas to determine the true prevalence of schistosome infection. The rMEA-IgG-ELISA results suggest that this assay can be valuable when used as a screening tool during epidemiological surveys followed by more specific assays as a robust parasitological evaluation. To overcome the complexity of ELISA in the field, a second-generation of antibody-based RDTs has already been proposed, as well as the detection of antigen together in a multiplex strip on a reader [60]. In this study, we have demonstrated this initial step successfully.

## Acknowledgments

We would like to thank the people from the rural communities of Montes Claros, Minas Gerais for their collaboration and the warm reception during the field activities. We are grateful for the excellent cooperation with the technical team of the Zoonosis Control Center, Montes Claros, MG, Brazil. Special thanks to Nuria Negrao and Flora Kano for teachings about recombinant proteins. Also, our thanks to Jessica Ramadhin for English review.

## References

1. GBD. Global, regional, and national age-sex specific mortality for 264 causes of death, 1980-2016: a systematic analysis for the Global Burden of Disease Study 2016. Lancet. 2017;390(10100):1151–210. doi: 10.1016/S0140-6736(17)32152-9. PubMed PMID: 28919116; PubMed Central PMCID: PMC5605883.

2. Steinmann P, Keiser J, Bos R, Tanner M, Utzinger J. Schistosomiasis and water resources development: systematic review, meta-analysis, and estimates of people at risk. Lancet Infect Dis. 2006;6(7):411–25. Epub 2006/06/23. doi: 10.1016/S1473-3099(06)70521-7. PubMed PMID: 16790382.

3. WHO. Accelerating work to overcome the global impact of neglected tropical diseases. A roadmap for implementation. 2012 [cited 2017 11/24]. Available from: http://www.who.int/neglected_diseases/NTD_RoadMap_2012_Fullversion.pdf?ua=1.

4. WHO/PAHO. Neglected infectious diseases in the Americas: Success stories and innovation to reach the neediest. 2016 [cited 2017 11/24]. Available from: http://www.paho.org/neglected-infectious-diseases-stories/http://iris.paho.org/xmlui/handle/123456789/31250.

5. WHO. Progress Report 2001–2011 and Strategic Plan 2012–2020 Geneva2013 [cited 2017 11/24]. Available from: http://www.who.int/iris/handle/10665/78074.

6. Katz N. Inquérito Nacional de Prevalência da Esquistossomose mansoni e Geo-helmintoses (2010-2015). Belo Horizonte: Instituto Rene Rachou - Fiocruz, 2018 March, 2018. Report No.: Contract No.: k197.

7. WHO/PAHO. Schistosomiasis Regional Meeting. Defining a road map toward verification of elimination of schistosomiasis transmission in Latin America and the Caribbean by 2020 2014 [cited 2017 11/24]. Available from: http://www.paho.org/hq/index.php?option=com_docman&task=doc_download&Itemid=270&gid=28841&lang=en.

8. Favre TC, Pereira AP, Beck LC, Galvao AF, Pieri OS. School-based and community-based actions for scaling-up diagnosis and treatment of schistosomiasis toward its elimination in an endemic area of Brazil. Acta Trop. 2015;149:155–62. Epub 2015/05/06. doi: 10.1016/j.actatropica.2015.04.024. PubMed PMID: 25940353.

9. Katz N, Chaves A, Pellegrino J. A simple device for quantitative stool thick-smear technique in Schistosomiasis mansoni. Rev Inst Med Trop Sao Paulo. 1972;14(6):397–400. Epub 1972/11/01. PubMed PMID: 4675644.

10. Siqueira LM, Gomes LI, Oliveira E, Oliveira ER, Oliveira AA, Enk MJ, et al. Evaluation of parasitological and molecular techniques for the diagnosis and assessment of cure of schistosomiasis mansoni in a low transmission area. Mem Inst Oswaldo Cruz. 2015;110(2):209–14. Epub 2015/05/07. doi: 10.1590/0074-02760140375. PubMed PMID: 25946244; PubMed Central PMCID: PMCPMC4489451.

11. Oliveira WJ, Magalhaes FDC, Elias AMS, de Castro VN, Favero V, Lindholz CG, et al. Evaluation of diagnostic methods for the detection of intestinal schistosomiasis in endemic areas with low parasite loads: Saline gradient, Helmintex, Kato-Katz and rapid urine test. PLoS Negl Trop Dis. 2018;12(2):e0006232. doi: 10.1371/journal.pntd.0006232. PubMed PMID: 29470516; PubMed Central PMCID: PMC5823366.

12. Siqueira LM, Couto FF, Taboada D, Oliveira AA, Carneiro NF, Oliveira E, et al. Performance of POC-CCA(R) in diagnosis of schistosomiasis mansoni in individuals with low parasite burden. Rev Soc Bras Med Trop. 2016;49(3):341–7. Epub 2016/07/08. doi: 10.1590/0037-8682-0070-2016. PubMed PMID: 27384831.

13. Espirito-Santo MC, Alvarado-Mora MV, Pinto PL, Sanchez MC, Dias-Neto E, Castilho VL, et al. Comparative Study of the Accuracy of Different Techniques for the Laboratory Diagnosis of Schistosomiasis Mansoni in Areas of Low Endemicity in Barra Mansa City, Rio de Janeiro State, Brazil. BioMed research international. 2015;2015:135689. doi: 10.1155/2015/135689. PubMed PMID: 26504777; PubMed Central PMCID: PMC4609343.

14. Pinheiro MC, Carneiro TR, Hanemann AL, Oliveira SM, Bezerra FS. The combination of three faecal parasitological methods to improve the diagnosis of schistosomiasis mansoni in a low endemic setting in the state of Ceara, Brazil. Mem Inst Oswaldo Cruz. 2012;107(7):873–6. PubMed PMID: 23147142.

15. Knopp S, Becker SL, Ingram KJ, Keiser J, Utzinger J. Diagnosis and treatment of schistosomiasis in children in the era of intensified control. Expert Rev Anti Infect Ther. 2013;11(11):1237–58. Epub 2013/10/17. doi: 10.1586/14787210.2013.844066. PubMed PMID: 24127662.

16. NTDs UtC. London Declaration on Neglected Tropical Diseases 2012 [cited 2017 11/24]. Available from: http://unitingtocombatntds.org/sites/default/files/resource_file/london_declaration_on_ntds.pdf.

17. WHO. WHA65.21. Elimination of schistosomiasis. Sixty-Fifth World Health Assembly; 21–26 May, 2012; Geneva, Switzerland: WHO; 2012. p. 36-7.

18. Gomes LI, Dos Santos Marques LH, Enk MJ, de Oliveira MC, Coelho PM, Rabello A. Development and evaluation of a sensitive PCR-ELISA system for detection of schistosoma infection in feces. PLoS Negl Trop Dis. 2010;4(4):e664. doi: 10.1371/journal.pntd.0000664. PubMed PMID: 20421918; PubMed Central PMCID: PMC2857640.

19. da Frota SM, Carneiro TR, Queiroz JA, Alencar LM, Heukelbach J, Bezerra FS. Combination of Kato-Katz faecal examinations and ELISA to improve accuracy of diagnosis of intestinal schistosomiasis in a low-endemic setting in Brazil. Acta Trop. 2011;120 Suppl 1:S138–41. doi: 10.1016/j.actatropica.2010.05.007. PubMed PMID: 20522322.

20. Silveira AM, Costa EG, Ray D, Suzuki BM, Hsieh MH, Fraga LA, et al. Evaluation of the CCA Immuno-Chromatographic Test to Diagnose Schistosoma mansoni in Minas Gerais State, Brazil. PLoS Negl Trop Dis. 2016;10(1):e0004357. Epub 2016/01/12. doi: 10.1371/journal.pntd.0004357. PubMed PMID: 26752073; PubMed Central PMCID: PMCPMC4709075.

21. van Dam GJ, Wichers JH, Ferreira TM, Ghati D, van Amerongen A, Deelder AM. Diagnosis of schistosomiasis by reagent strip test for detection of circulating cathodic antigen. J Clin Microbiol. 2004;42(12):5458–61. Epub 2004/12/08. doi: 10.1128/JCM.42.12.5458-5461.2004. PubMed PMID: 15583265; PubMed Central PMCID: PMCPMC535219.

22. Colley DG, Binder S, Campbell C, King CH, Tchuem Tchuente LA, N’Goran EK, et al. A five-country evaluation of a point-of-care circulating cathodic antigen urine assay for the prevalence of Schistosoma mansoni. Am J Trop Med Hyg. 2013;88(3):426–32. Epub 2013/01/23. doi: 10.4269/ajtmh.12-0639. PubMed PMID: 23339198; PubMed Central PMCID: PMCPMC3592520.

23. Tchuem Tchuente LA, Kuete Fouodo CJ, Kamwa Ngassam RI, Sumo L, Dongmo Noumedem C, Kenfack CM, et al. Evaluation of circulating cathodic antigen (CCA) urine-tests for diagnosis of Schistosoma mansoni infection in Cameroon. PLoS Negl Trop Dis. 2012;6(7):e1758. Epub 2012/08/04. doi: 10.1371/journal.pntd.0001758. PubMed PMID: 22860148; PubMed Central PMCID: PMCPMC3409114.

24. Shane HL, Verani JR, Abudho B, Montgomery SP, Blackstock AJ, Mwinzi PN, et al. Evaluation of urine CCA assays for detection of Schistosoma mansoni infection in Western Kenya. PLoS Negl Trop Dis. 2011;5(1):e951. doi: 10.1371/journal.pntd.0000951. PubMed PMID: 21283613; PubMed Central PMCID: PMC3026766.

25. Coulibaly JT, Knopp S, N’Guessan NA, Silue KD, Furst T, Lohourignon LK, et al. Accuracy of urine circulating cathodic antigen (CCA) test for Schistosoma mansoni diagnosis in different settings of Cote d’Ivoire. PLoS Negl Trop Dis. 2011;5(11):e1384. Epub 2011/12/02. doi: 10.1371/journal.pntd.0001384. PubMed PMID: 22132246; PubMed Central PMCID: PMCPMC3222626.

26. Kittur N, Castleman JD, Campbell CH, Jr., King CH, Colley DG. Comparison of Schistosoma mansoni Prevalence and Intensity of Infection, as Determined by the Circulating Cathodic Antigen Urine Assay or by the Kato-Katz Fecal Assay: A Systematic Review. Am J Trop Med Hyg. 2016;94(3):605–10. Epub 2016/01/13. doi: 10.4269/ajtmh.15-0725. PubMed PMID: 26755565; PubMed Central PMCID: PMCPMC4775897.

27. Grenfell RFQ, Taboada D, Coutinho LA, Pedrosa MLC, Assis JV, Oliveira MSP, et al. Innovative methodology for point-of-care circulating cathodic antigen with rapid urine concentration for use in the field for detecting low Schistosoma mansoni infection and for control of cure with high accuracy. Trans R Soc Trop Med Hyg. 2018. doi: 10.1093/trstmh/try014. PubMed PMID: 29522211.

28. Coelho PM, Siqueira LM, Grenfell RF, Almeida NB, Katz N, Almeida A, et al. Improvement of POC-CCA Interpretation by Using Lyophilization of Urine from Patients with Schistosoma mansoni Low Worm Burden: Towards an Elimination of Doubts about the Concept of Trace. PLoS Negl Trop Dis. 2016;10(6):e0004778. Epub 2016/06/22. doi: 10.1371/journal.pntd.0004778. PubMed PMID: 27326453; PubMed Central PMCID: PMCPMC4915691.

29. Silva JDDF, Pinheiro MCC, Sousa MS, Gomes VDS, Castro IMN, Ramos ANJ, et al. Detection of schistosomiasis in an area directly affected by the Sao Francisco River large-scale water transposition project in the Northeast of Brazil. Rev Soc Bras Med Trop. 2017;50(5):658–65. doi: 10.1590/0037-8682-0299-2017. PubMed PMID: 29160513.

30. Bezerra FSM, Leal JKF, Sousa MS, Pinheiro MCC, Ramos AN, Jr., Silva-Moraes V, et al. Evaluating a Point-of-Care Circulating Cathodic Antigen test (POC-CCA) to detect Schistosoma mansoni infections in a low endemic area in north-eastern Brazil. Acta Trop. 2018. doi: 10.1016/j.actatropica.2018.03.002. PubMed PMID: 29526480.

31. Lindholz CG, Favero V, Verissimo CM, Candido RRF, de Souza RP, Dos Santos RR, et al. Study of diagnostic accuracy of Helmintex, Kato-Katz, and POC-CCA methods for diagnosing intestinal schistosomiasis in Candeal, a low intensity transmission area in northeastern Brazil. PLoS Negl Trop Dis. 2018;12(3):e0006274. doi: 10.1371/journal.pntd.0006274. PubMed PMID: 29518081; PubMed Central PMCID: PMC5843168.

32. Ferreira FT, Fidelis TA, Pereira TA, Otoni A, Queiroz LC, Amancio FF, et al. Sensitivity and specificity of the circulating cathodic antigen rapid urine test in the diagnosis of Schistosomiasis mansoni infection and evaluation of morbidity in a low-endemic area in Brazil. Rev Soc Bras Med Trop. 2017;50(3):358–64. Epub 2017/07/13. doi: 10.1590/0037-8682-0423-2016. PubMed PMID: 28700054.

33. Marinho CC, Groberio AC, Silva C, Lima TLF, Santos RCD, Araujo LG, et al. Morbidity of schistosomiasis mansoni in a low endemic setting in Ouro Preto, Minas Gerais, Brazil. Rev Soc Bras Med Trop. 2017;50(6):805–11. doi: 10.1590/0037-8682-0009-2017. PubMed PMID: 29340458.

34. Espirito-Santo MC, Sanchez MC, Sanchez AR, Alvarado-Mora MV, Castilho VL, Goncalves EM, et al. Evaluation of the sensitivity of IgG and IgM ELISA in detecting Schistosoma mansoni infections in a low endemicity setting. Eur J Clin Microbiol Infect Dis. 2014;33(12):2275–84. Epub 2014/07/18. doi: 10.1007/s10096-014-2196-6. PubMed PMID: 25030291.

35. Grenfell RF, Martins W, Enk M, Almeida A, Siqueira L, Silva-Moraes V, et al. Schistosoma mansoni in a low-prevalence area in Brazil: the importance of additional methods for the diagnosis of hard-to-detect individual carriers by low-cost immunological assays. Mem Inst Oswaldo Cruz. 2013;108(3). Epub 2013/06/20. doi: 10.1590/S0074-02762013000300011. PubMed PMID: 23778663; PubMed Central PMCID: PMCPMC4005562.

36. Goncalves MM, Barreto MG, Peralta RH, Gargioni C, Goncalves T, Igreja RP, et al. Immunoassays as an auxiliary tool for the serodiagnosis of Schistosoma mansoni infection in individuals with low intensity of egg elimination. Acta Trop. 2006;100(1-2):24–30. Epub 2006/10/31. doi: 10.1016/j.actatropica.2006.09.004. PubMed PMID: 17069742.

37. Carneiro TR, Pinheiro MC, de Oliveira SM, Hanemann AL, Queiroz JA, Bezerra FS. Increased detection of schistosomiasis with Kato-Katz and SWAP-IgG-ELISA in a Northeastern Brazil low-intensity transmission area. Rev Soc Bras Med Trop. 2012;45(4):510–3. PubMed PMID: 22930048.

38. Oliveira EJ, Kanamura HY, Lima DM. Efficacy of an enzyme-linked immunosorbent assay as a diagnostic tool for schistosomiasis mansoni in individuals with low worm burden. Mem Inst Oswaldo Cruz. 2005;100(4):421–5. doi: /S0074-02762005000400013. PubMed PMID: 16113891.

39. Alarcon de Noya B, Ruiz R, Losada S, Colmenares C, Contreras R, Cesari IM, et al. Detection of schistosomiasis cases in low-transmission areas based on coprologic and serologic criteria The Venezuelan experience. Acta Trop. 2007;103(1):41–9. doi: 10.1016/j.actatropica.2007.04.018. PubMed PMID: 17606217.

40. Lambertucci JR, Drummond SC, Voieta I, de Queiroz LC, Pereira PP, Chaves BA, et al. An outbreak of acute Schistosoma mansoni Schistosomiasis in a nonendemic area of Brazil: a report on 50 cases, including 5 with severe clinical manifestations. Clin Infect Dis. 2013;57(1):e1–6. Epub 2013/03/28. doi: 10.1093/cid/cit157. PubMed PMID: 23532472.

41. Grenfell RF, Martins W, Drummond SC, Antunes CM, Voieta I, Otoni A, et al. Acute schistosomiasis diagnosis: a new tool for the diagnosis of schistosomiasis in a group of travelers recently infected in a new focus of Schistosoma mansoni. Rev Soc Bras Med Trop. 2013;46(2):208–13. Epub 2013/06/07. doi: 10.1590/0037-8682-0064-2012. PubMed PMID: 23740077.

42. Kinkel HF, Dittrich S, Baumer B, Weitzel T. Evaluation of eight serological tests for diagnosis of imported schistosomiasis. Clin Vaccine Immunol. 2012;19(6):948–53. Epub 2012/03/24. doi: 10.1128/CVI.05680-11. PubMed PMID: 22441394; PubMed Central PMCID: PMCPMC3370443.

43. Doenhoff MJ, Chiodini PL, Hamilton JV. Specific and sensitive diagnosis of schistosome infection: can it be done with antibodies? Trends Parasitol. 2004;20(1):35–9. Epub 2004/01/01. PubMed PMID: 14700588.

44. Alarcon de Noya B, Colmenares C, Lanz H, Caracciolo MA, Losada S, Noya O. Schistosoma mansoni: immunodiagnosis is improved by sodium metaperiodate which reduces cross-reactivity due to glycosylated epitopes of soluble egg antigen. Exp Parasitol. 2000;95(2):106–12. doi: 10.1006/expr.2000.4515. PubMed PMID: 10910711.

45. Seliger B, Kellner R. Design of proteome-based studies in combination with serology for the identification of biomarkers and novel targets. Proteomics. 2002;2(12):1641–51. doi: 10.1002/1615-9861(200212)2:12<1641::AID-PROT1641>3.0.CO;2-B. PubMed PMID: 12469333.

46. Coelho VT, Oliveira JS, Valadares DG, Chavez-Fumagalli MA, Duarte MC, Lage PS, et al. Identification of proteins in promastigote and amastigote-like Leishmania using an immunoproteomic approach. PLoS Negl Trop Dis. 2012;6(1):e1430. doi: 10.1371/journal.pntd.0001430. PubMed PMID: 22272364; PubMed Central PMCID: PMC3260309.

47. Zhang M, Fu Z, Li C, Han Y, Cao X, Han H, et al. Screening diagnostic candidates for schistosomiasis from tegument proteins of adult Schistosoma japonicum using an immunoproteomic approach. PLoS Negl Trop Dis. 2015;9(2):e0003454. doi: 10.1371/journal.pntd.0003454. PubMed PMID: 25706299; PubMed Central PMCID: PMC4338221.

48. Losada S, Sabatier L, Hammann P, Guillier C, Matos C, Bermudez H, et al. A combined proteomic and immunologic approach for the analysis of Schistosoma mansoni cercariae and adult worm protein extracts and the detection of one of the vaccine candidates, Sm28GST, from a Venezuelan parasite isolate. Investigacion clinica. 2011;52(2):121–39. PubMed PMID: 21866785.

49. Zhong ZR, Zhou HB, Li XY, Luo QL, Song XR, Wang W, et al. Serological proteome-oriented screening and application of antigens for the diagnosis of Schistosomiasis japonica. Acta Trop. 2010;116(1):1–8. doi: 10.1016/j.actatropica.2010.04.014. PubMed PMID: 20451489.

50. Wilson RA, Langermans JA, van Dam GJ, Vervenne RA, Hall SL, Borges WC, et al. Elimination of Schistosoma mansoni Adult Worms by Rhesus Macaques: Basis for a Therapeutic Vaccine? PLoS Negl Trop Dis. 2008;2(9):e290. doi: 10.1371/journal.pntd.0000290. PubMed PMID: 18820739; PubMed Central PMCID: PMC2553480.

51. Mutapi F, Burchmore R, Mduluza T, Foucher A, Harcus Y, Nicoll G, et al. Praziquantel treatment of individuals exposed to Schistosoma haematobium enhances serological recognition of defined parasite antigens. J Infect Dis. 2005;192(6):1108–18. doi: 10.1086/432553. PubMed PMID: 16107967.

52. Ludolf F, Patrocinio PR, Correa-Oliveira R, Gazzinelli A, Falcone FH, Teixeira-Ferreira A, et al. Serological screening of the Schistosoma mansoni adult worm proteome. PLoS Negl Trop Dis. 2014;8(3):e2745. doi: 10.1371/journal.pntd.0002745. PubMed PMID: 24651847; PubMed Central PMCID: PMC3961189.

53. Smith H, Doenhoff M, Aitken C, Bailey W, Ji M, Dawson E, et al. Comparison of Schistosoma mansoni soluble cercarial antigens and soluble egg antigens for serodiagnosing schistosome infections. PLoS Negl Trop Dis. 2012;6(9):e1815. doi: 10.1371/journal.pntd.0001815. PubMed PMID: 23029577; PubMed Central PMCID: PMC3441401.

54. Meevissen MH, Wuhrer M, Doenhoff MJ, Schramm G, Haas H, Deelder AM, et al. Structural characterization of glycans on omega-1, a major Schistosoma mansoni egg glycoprotein that drives Th2 responses. Journal of proteome research. 2010;9(5):2630–42. doi: 10.1021/pr100081c. PubMed PMID: 20178377.

55. Coelho PM, Jurberg AD, Oliveira AA, Katz N. Use of a saline gradient for the diagnosis of schistosomiasis. Mem Inst Oswaldo Cruz. 2009;104(5):720–3. Epub 2009/10/13. PubMed PMID: 19820832.

56. Ashton PD, Harrop R, Shah B, Wilson RA. The schistosome egg: development and secretions. Parasitology. 2001;122(Pt 3):329–38. PubMed PMID: 11289069.

57. Woodward MP, Young WW, Jr., Bloodgood RA. Detection of monoclonal antibodies specific for carbohydrate epitopes using periodate oxidation. Journal of immunological methods. 1985;78(1):143–53. PubMed PMID: 2580026.

58. Invitrogen. A universal technology to clone DNA sequences for functional analysis and expression in multiple systems. 2003.

59. Greiner M. Two-graph receiver operating characteristic (TG-ROC): a Microsoft-EXCEL template for the selection of cut-off values in diagnostic tests. Journal of immunological methods. 1995;185(1):145–6. PubMed PMID: 7665897.

60. Corstjens PL, De Dood CJ, Kornelis D, Fat EM, Wilson RA, Kariuki TM, et al. Tools for diagnosis, monitoring and screening of Schistosoma infections utilizing lateral-flow based assays and upconverting phosphor labels. Parasitology. 2014;141(14):1841–55. Epub 2014/06/17. doi: 10.1017/S0031182014000626. PubMed PMID: 24932595; PubMed Central PMCID: PMCPMC4265670.

61. Colley DG, Andros TS, Campbell CH, Jr. Schistosomiasis is more prevalent than previously thought: what does it mean for public health goals, policies, strategies, guidelines and intervention programs? Infect Dis Poverty. 2017;6(1):63. Epub 2017/03/23. doi: 10.1186/s40249-017-0275-5. PubMed PMID: 28327187; PubMed Central PMCID: PMCPMC5361841.

62. Igreja RP, Matos JA, Goncalves MM, Barreto MM, Peralta JM. Schistosoma mansoni-related morbidity in a low-prevalence area of Brazil: a comparison between egg excretors and seropositive non-excretors. Ann Trop Med Parasitol. 2007;101(7):575–84. Epub 2007/09/20. doi: 10.1179/136485907X229086. PubMed PMID: 17877876.

63. Makarova E, Goes TS, Leite MF, Goes AM. Detection of IgG binding to Schistosoma mansoni recombinant protein RP26 is a sensitive and specific method for acute schistosomiasis diagnosis. Parasitol Int. 2005;54(1):69–74. doi: 10.1016/j.parint.2004.12.001. PubMed PMID: 15710554.

64. Tanigawa C, Fujii Y, Miura M, Nzou SM, Mwangi AW, Nagi S, et al. Species-Specific Serological Detection for Schistosomiasis by Serine Protease Inhibitor (SERPIN) in Multiplex Assay. PLoS Negl Trop Dis. 2015;9(8):e0004021. doi: 10.1371/journal.pntd.0004021. PubMed PMID: 26291988; PubMed Central PMCID: PMC4546333.

65. Beck L, Van-Lume DS, Souza JR, Morais CN, Melo WG, Xavier E, et al. Evaluation of tests based on the antibody response to keyhole limpet haemocyanin and soluble egg antigen to differentiate acute and chronic human schistosomiasis mansoni. Mem Inst Oswaldo Cruz. 2004;99(5 Suppl 1):97–8. doi: /S0074-02762004000900017. PubMed PMID: 15486643.

66. de Assis RR, Ludolf F, Nakajima R, Jasinskas A, Oliveira GC, Felgner PL, et al. A next-generation proteome array for Schistosoma mansoni. International journal for parasitology. 2016;46(7):411–5. doi: 10.1016/j.ijpara.2016.04.001. PubMed PMID: 27131510.

67. Mathieson W, Wilson RA. A comparative proteomic study of the undeveloped and developed Schistosoma mansoni egg and its contents: the miracidium, hatch fluid and secretions. International journal for parasitology. 2010;40(5):617–28. doi: 10.1016/j.ijpara.2009.10.014. PubMed PMID: 19917288.

68. Curwen RS, Ashton PD, Johnston DA, Wilson RA. The Schistosoma mansoni soluble proteome: a comparison across four life-cycle stages. Molecular and biochemical parasitology. 2004;138(1):57–66. doi: 10.1016/j.molbiopara.2004.06.016. PubMed PMID: 15500916.

69. Cass CL, Johnson JR, Califf LL, Xu T, Hernandez HJ, Stadecker MJ, et al. Proteomic analysis of Schistosoma mansoni egg secretions. Molecular and biochemical parasitology. 2007;155(2):84–93. doi: 10.1016/j.molbiopara.2007.06.002. PubMed PMID: 17644200; PubMed Central PMCID: PMC2077830.

70. van Balkom BW, van Gestel RA, Brouwers JF, Krijgsveld J, Tielens AG, Heck AJ, et al. Mass spectrometric analysis of the Schistosoma mansoni tegumental sub-proteome. Journal of proteome research. 2005;4(3):958–66. doi: 10.1021/pr050036w. PubMed PMID: 15952743.

71. Cao M, Chao H, Doughty BL. Cloning of a cDNA encoding an egg antigen homologue from Schistosoma mansoni. Molecular and biochemical parasitology. 1993;58(1):169–71. PubMed PMID: 8459829.

72. Abouel-Nour MF, Lotfy M, Attallah AM, Doughty BL. Schistosoma mansoni major egg antigen Smp40: molecular modeling and potential immunoreactivity for anti-pathology vaccine development. Mem Inst Oswaldo Cruz. 2006;101(4):365–72. PubMed PMID: 16951805.

73. Nene V, Dunne DW, Johnson KS, Taylor DW, Cordingley JS. Sequence and expression of a major egg antigen from Schistosoma mansoni. Homologies to heat shock proteins and alpha-crystallins. Molecular and biochemical parasitology. 1986;21(2):179–88. PubMed PMID: 3097539.

74. Cavalcanti MG, Silva LF, Peralta RH, Barreto MG, Peralta JM. Schistosomiasis in areas of low endemicity: a new era in diagnosis. Trends Parasitol. 2013;29(2):75–82. Epub 2013/01/08. doi: 10.1016/j.pt.2012.11.003. PubMed PMID: 23290589.

75. Carvalho GB, Pacifico LG, Pimenta DL, Siqueira LM, Teixeira-Carvalho A, Coelho PM, et al. Evaluation of the use of C-terminal part of the Schistosoma mansoni 200kDa tegumental protein in schistosomiasis diagnosis and vaccine formulation. Exp Parasitol. 2014;139:24–32. Epub 2014/02/25. doi: 10.1016/j.exppara.2014.02.003. PubMed PMID: 24560833.

76. Grenfell RF, Silva-Moraes V, Taboada D, de Mattos AC, de Castro AK, Coelho PM. Immunodiagnostic methods: what is their role in areas of low endemicity? ScientificWorldJournal. 2012;2012:593947. Epub 2013/01/16. doi: 10.1100/2012/593947. PubMed PMID: 23319886; PubMed Central PMCID: PMCPMC3539347.

77. Kalenda YD, Kato K, Goto Y, Fujii Y, Hamano S. Tandem repeat recombinant proteins as potential antigens for the sero-diagnosis of Schistosoma mansoni infection. Parasitol Int. 2015;64(6):503–12. doi: 10.1016/j.parint.2015.06.012. PubMed PMID: 26148816.

78. Sarhan RM, Aminou HA, Saad GA, Ahmed OA. Comparative analysis of the diagnostic performance of adult, cercarial and egg antigens assessed by ELISA, in the diagnosis of chronic human Schistosoma mansoni infection. Parasitology research. 2014;113(9):3467–76. doi: 10.1007/s00436-014-4017-3. PubMed PMID: 25028207.

79. Grenfell R, Harn DA, Tundup S, Da’dara A, Siqueira L, Coelho PM. New approaches with different types of circulating cathodic antigen for the diagnosis of patients with low Schistosoma mansoni load. PLoS Negl Trop Dis. 2013;7(2):e2054. Epub 2013/03/08. doi: 10.1371/journal.pntd.0002054. PubMed PMID: 23469295; PubMed Central PMCID: PMCPMC3585039.

80. El Aswad Bel D, Doenhoff MJ, El Hadidi AS, Schwaeble WJ, Lynch NJ. Use of recombinant calreticulin and cercarial transformation fluid (CTF) in the serodiagnosis of Schistosoma mansoni. Immunobiology. 2011;216(3):379–85. Epub 2010/08/10. doi: 10.1016/j.imbio.2010.06.014. PubMed PMID: 20691496.

81. Dawson EM, Sousa-Figueiredo JC, Kabatereine NB, Doenhoff MJ, Stothard JR. Intestinal schistosomiasis in pre school-aged children of Lake Albert, Uganda: diagnostic accuracy of a rapid test for detection of anti-schistosome antibodies. Trans R Soc Trop Med Hyg. 2013;107(10):639–47. Epub 2013/08/27. doi: 10.1093/trstmh/trt077. PubMed PMID: 23976783.

82. Sorgho H, Bahgat M, Poda JN, Song W, Kirsten C, Doenhoff MJ, et al. Serodiagnosis of Schistosoma mansoni infections in an endemic area of Burkina Faso: performance of several immunological tests with different parasite antigens. Acta Trop. 2005;93(2):169–80. doi: 10.1016/j.actatropica.2004.10.006. PubMed PMID: 15652331.

